# Shaping the Glycan Landscape: Hidden relationships between linkage and ring distortions induced by carbohydrate-active enzymes

**DOI:** 10.1101/2025.06.27.661892

**Authors:** Isabell Louise Grothaus, Paul Spellerberg, Carme Rovira, Lucio Colombi Ciacchi

**Affiliations:** Hybrid Materials Interfaces Group, Bremen Center for Computational Materials Science and MAPEX Center for Materials and Processes, University of Bremen, 28359 Bremen, Germany; Malopolska Centre of Biotechnology, Jagiellonian University, 31-007 Krakow, Poland; Department of Theoretical Biophysics, Max Planck Institute for Biophysics, 60438 Frankfurt, Germany; Faculty for Biology and Chemistry, University of Bremen, 28359 Bremen, Germany; Departament de Química Inorgà nica i Orgà nica & IQTCUB, Universitat de Barcelona, Barcelona 08028, Spain; Institució Catalana de Recerca i Estudis Avançats (ICREA), Barcelona 08020, Spain

## Abstract

Carbohydrate-active enzymes (CAZymes) catalyze glycan remodeling by forming and cleaving glycosidic bonds in diverse biological environments. Often, a key aspect of their catalytic mechanism is a monosaccharide chair-to-boat distortion that brings the substrate from a stable solution conformation to a reactive state. Using enhanced-sampling molecular dynamics simulations, we demonstrate that the ring distortion experienced by the glycan M5G0 upon binding to the Golgi *α*-mannosidase II (MII) enzyme actively correlates with a change of its global conformation. In solution, M5G0 adopts diverse conformers, all favoring the ^4^*C*_1_ chair for the mannose at subsite *-1* with respect to the bond cleavage point. Binding to MII narrows the glycan’s phase space to only two low-energy conformers, which respectively correlate with the two distinct pucker states ^4^*C*_1_ and ^0^*H*_5_. Key factors driving this phase-space reshaping include binding to specific amino acids and a Zn^2+^ ion in the catalytic site. Comparative studies with ER *α*-mannosidase I show a different mechanism, where the enzyme enforces glycan conformations and ring distortion of the substrate independently. Our findings provide mechanistic insights into CAZyme specificity and effectiveness, laying the groundwork for the design of selective inhibitors targeting glycosylation-related diseases, including cancer.

## Introduction

Carbohydrate-active enzymes (CAZymes) account for 1 to 3 % of most genomes. They are responsible for the modification of the cell glycome through glycosidic bond breakage and formation, particularly by glycoside hydrolases (GH) and glycosyltransferases (GT), respectively. ^1^ The vast diversity and complexity of glycan substrates requires a wide portfolio of discrete folds, amounting to 189 GH and 137 GT families with specific carbohydrate-binding and reshaping activities.^2,a^ Glycans are comprised of sugar monomers (monosaccharides) assembled into a variety of branched structures through the formation of glycosidic linkages. Owing to the diversity of monosaccharides, linkage positions, orientations and degrees of branching, glycans inherit a much larger chemical and structural variety compared to nucleic acids and proteins. Their behavior is characterized by a high flexibility, which manifests itself in rapid transitions between individual conformers (Figure 1 A/B).^3^ These can be differentiated by the dihedral (torsional) angles *ϕ, ψ* and *ω* along the glycosidic bonds, and by the Cremer-Pople parameters *ϕ* and *θ* determining the puckering of the monosaccharide rings (Figure 1 C/D).^4^ The probability distributions of all dihedral and puckering degrees of freedom adopted by a glycan during its dynamics define its conformational phase space, which captures and quantifies its structural character and is associated with its biological functions.

**Figure 1.**
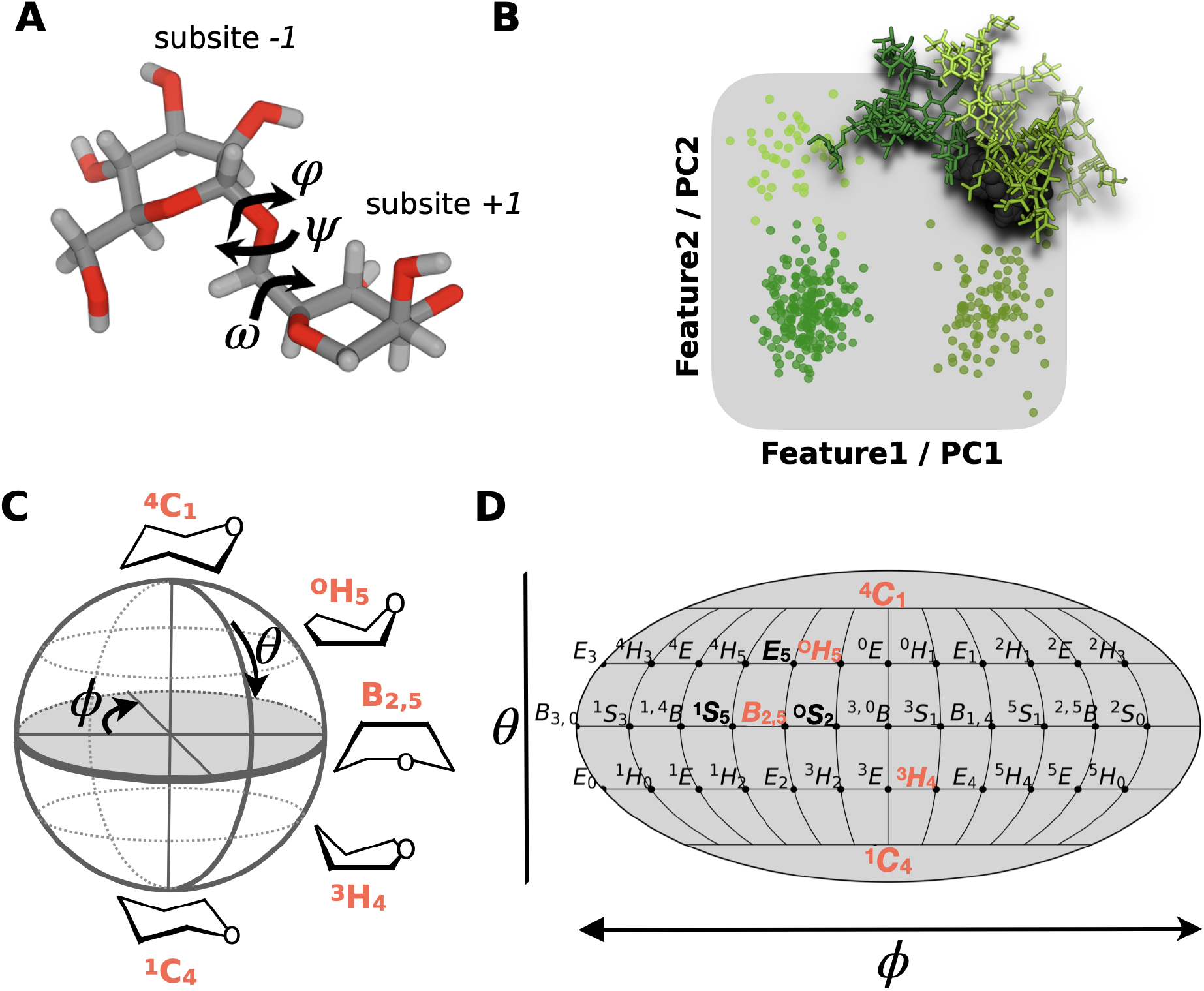
Representation of the glycan’s conformational phase space. **A** Glycans are flexible molecules, adopting various distinct conformers that are characterized by the dihedral angles *ϕ, ψ* and *ω* along the glycosidic bonds connecting the individual monosaccharides e.g. mannoses located at subsite *-1* and *+1* in a CAZyme binding site. The atomistic structure is represented in licorice style with oxygen atoms in red, hydrogens in white and carbons in gray. **B** The conformational phase space can be best represented in low dimensions employing features as axis that can be identified by e.g. principle component analysis. Points in this latent space represent individual structures that are colored by their respective conformer. Atomistic structures colored in green represent three different conformers of the glycan M5G0. **C** An additional degree of freedom is the flexibility of the pyranose ring, whose distortion is quantified by the Cremer-Pople pucker coordinates *θ* and *ϕ* and **D** plotted using a Mollweide projection, a pseudocylindrical map with equal-area, meaning that areas, densities and, thus, free energy values are preserved. Labels represent individual ring shapes. ^4^

CAZymes evolved a remarkable binding specificity for different glycans. Especially the cleaving mechanism of GH enzymes is particularly intriguing, because the glycan substrates very often undergo a ring distortion at the monosaccharide subsite *-1* with respect to the bond cleavage point upon binding to the catalytic site (Figure 2 A). ^5,6^ This puckering is necessary to achieve an axial linkage or oxocarbenium-ion (OCI) character, resulting in a higher catalytic efficiency for the adjacent bond cleavage, irrespective of whether the hydrolytic mechanism is inverting or retaining.^5,6^ The precise itinerary from the reactant state (also known as the Michaelis complex) to the transition and the product states along the catalytic reaction might vary for different substrates and GH families. However, ring distortions away from the most stable ring shapes ^4^*C*_1_ and ^1^*C*_4_ adopted in solution are likely to be observed in all these states. What causes the ring distortion, especially before chemical attack occurs, is still unclear. It is debated whether the residues of the catalytic site promote the ring distortion via a chemical action, such as strong polarization of the ring’s functional groups, ^5^ or whether the adoption of specific glycan conformations far from the solution equilibrium results in steric activation of otherwise inaccessible pucker states (Figure 2 A). Knowledge about the precise catalytic itinerary and its root-causes is relevant for i) a better understanding of CAZymes’ reaction mechanisms, ii) a prediction of substrate specificity, and iii) the design of inhibitors for therapeutic enzyme inhibitors. The latter are crucial in a wide field of applications such as antiviral medicines, treatment of genetic disorders, mitigation of diabetes, or cancer therapy. ^7,8^

**Figure 2.**
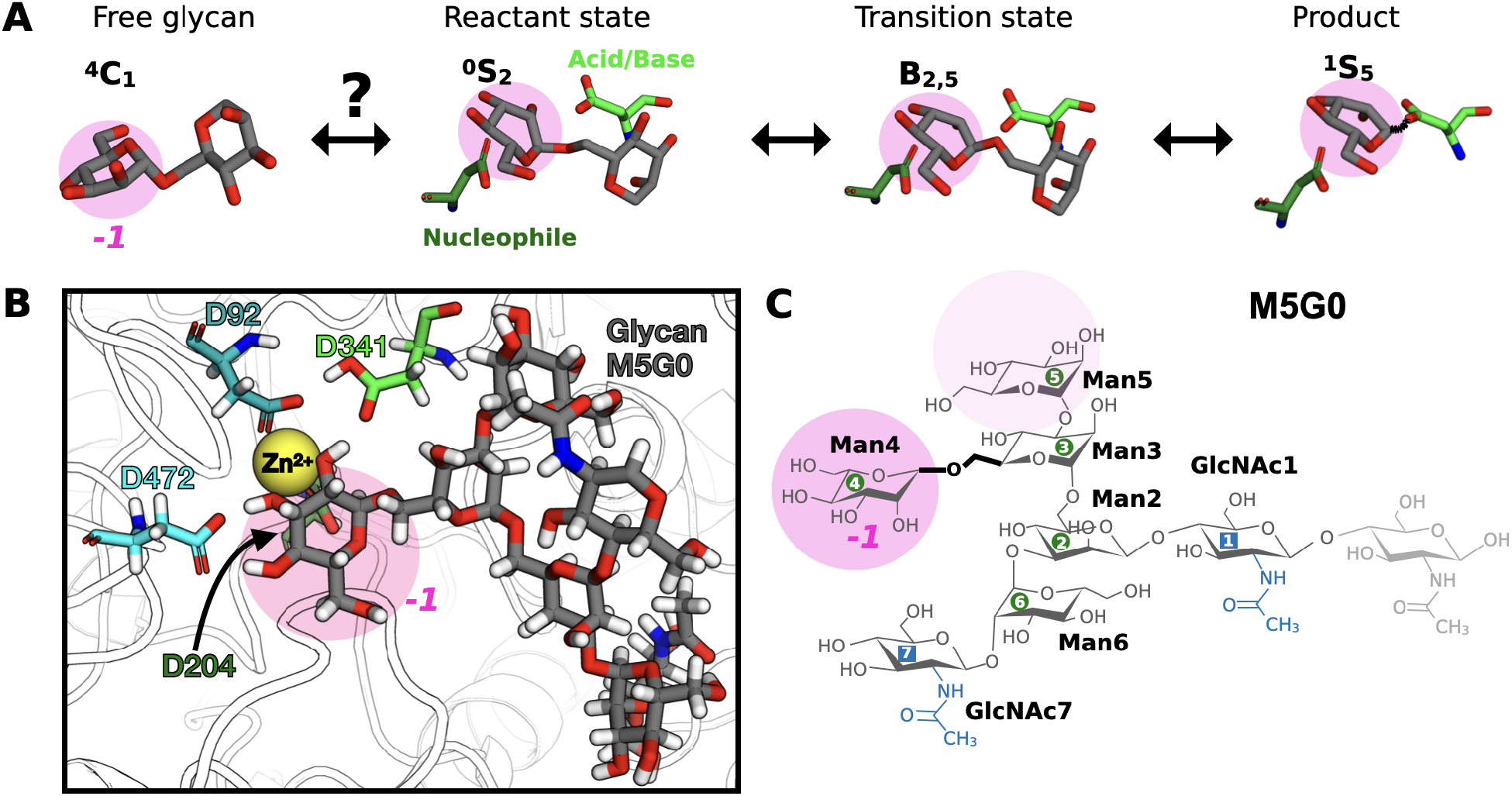
Catalytic details of Golgi *α*-mannosidase II. **A** The catalytic itinerary of the glycosylation step, depicting the transition from the solution equilibrium ^4^*C*_1_ chair to the product in the ^1^*S*_5_ skew-boat. The monosaccharide at subsite *-1* is the residue to be cleaved off. **B** The binding configuration of glycan M5G0 within the catalytic site, highlighting the Zn^2+^ ion and key amino acid residues (D92, D204, D341, and D472). Atoms are displayed in a licorice style: the glycan has gray carbon atoms, while amino acid carbons are color-coded by residue type. Oxygen atoms are red, nitrogen atoms are blue, and hydrogen atoms are white. **C** Atomistic structure and nomenclature of M5G0 with labeling of linkages according to neighboring residue numbers. The Man3-Man4 linkage is highlighted in bold and to be cleaved by MII. In all panels, the to-be-cleaved Man4, positioned at the catalytic subsite *-1*, is emphasized with a transparent pink circle.

In this work, we investigate the origins of ring distortion for glycan substrates of *α*-mannosidase I (MI) and *α*-mannosidase II (MII). Mannosidases make up almost 10 % of all GH families, and are crucial for the correct processing of protein N-glycosylation. ^9^ Especially MII is of great biological and medical relevance, enabling the transition from high-mannose type to complex N-glycans. The enzyme is known to be overexpressed in cancer types like colon, skin and breast cancer, resulting in altered glycoforms on cell membrane glycoproteins, which could be correlated with metastasis growth and disease progression.^10–12^ Indeed, inhibition of MII minimized the formation of complex N-glycans and was associated with reduced tumor growth and metastasis.^13^

MII catalyzes the reduction of its substrate M5G0 by cleaving the *α*1-6 linkage between the mannose (Man) Man3 and Man4, resulting in an intermediate substrate, which is eventually processed to a complex glycan after a further reduction step (Figure 2 C). Studies based on X-ray diffraction and QM/MM calculations have established that, starting from its equilibrium chair structure in solution, Man4 transitions through various boat states in the enzyme binding pocket during the enzymatic cleavage. The shapes so far identified along the pucker itinerary are: ^4^*C*_1_ *→*^*O*^ *S*_2_*/B*_2,5_ *→ B*_2,5_ *→*^1^ *S*_5_.^14,15^ The oxocarbenium ion-like transition state is a prerequisite for hydrolysis, because the axial orientation of the 2-hydroxyl group in a ^4^*C*_1_ chair is unfavourable for the required nucleophilic substitution reaction at the anomeric position.^9^ The ring distortion seems to be assisted by a Zn^2+^ ion in the binding pocket, which binds to the hydroxyl groups of Man4 and could possibly alter the monosaccharide ring shape.^14,16^ Analogously, the inverting *α*-mannosidase I (MI) catalyses the reduction of its substrate M9 by cleaving a terminal mannose transitioning from its equilibrium ^1^*C*_4_ chair to a ^3^*H*_4_ half-chair.^17^ In this case, the reaction is assisted by a Ca^2+^ ion in the enzyme’s binding pocket.

Current experimental insights into the pucker itineraries followed by mannosidase substrates are limited to static intermediates stabilized in mutant crystal structures. For this reason, the factors influencing the dynamics leading to the distorted ring shape of the initial reactant state remain mysterious.^5^ Computational studies based on all-atom molecular dynamics (MD) simulations have the potential to shed light onto this issue, given that popular force-field parametrizations have been shown to accurately reproduce ring distortions across different states of the catalytic itinerary.^14^ However, previous studies could not explore the entire conformational phase spaces of the glycan substrates due to limited receptor and ligand flexibility in molecular docking algorithms, and the short timescales accessible by quantum mechanics/molecular mechanics (QM/MM) approaches or unbiased MD simulations.^18^ For this reason, here we employ enhanced-sampling MD simulations that give justice to the dynamical flexibility of glycans, utilizing a previously introduced REST-RECT methodology that ensures comprehensive sampling of their conformational phase spaces. ^3^ This will allow us to quantify the significant phase-space reshaping undertaken by M5G0 and M9 upon binding to the catalytic sites of MII and MI, and to identify the several key contributors to the peculiar ring distortions of the reactant states.

## Methods

### Setup of simulation systems

All-atom MD simulations were performed for the native or mutated CAZymes MII (PDB 3CZN) and MI (PDB 5KIJ) with their co-crystallized natural substrates M5G0 and M9. Initial structures and force field parameters were generated via CHARMM-GUI ^19,20^, setting the pH to 7.0 and preserving the glycan structures and components. This means that in both cases the first GlcNAc residue at the non-reducing end, which was not resolved in the crystal structures, is not included in the models. For MII, all amino acids 31 - 1044 were included in the model. The residue D341 was protonated, as known for the active enzyme, and the mutation D204A used to stabilize the crystal structure was reversed to a deprotonated aspartic acid. In order to construct the modified substrate M5 in the binding site of MII, GlcNAc7 was deleted from the M5G0 structure and replaced by an OH group. For MI, all amino acids 245 - 696 were included in the model. The La^3+^ ion used experimentally for resolving the crystal structure from X-ray data was converted to a Ca^2+^ ion, as in the natural enzyme. The atomic coordinates of the water molecules bound to the ion were preserved, because the ion hydration shell is crucial for its correct positioning in the catalytic site. Residues E602, E663, E689, D255 and H385 of MI were protonated and a disulfide bond introduced between residues 527 and 556. All systems were solvated using a 10 Å thick water layer in either cubic or dodecahedron boxes and neutralized using K^+^ or Cl^*−*^ ions, as required. Simulation setups of free glycans in solution were also constructed via CHARMM-GUI’s Glycan Modeller, based on averaged structures from the Glycan Fragment Database^19,21–23^ and solvated and neutralized as described above.

Enhanced-sampling MD simulations were performed with GROMACS 2022^24^, patched with the PLUMED package version 2.8 ^25^. The Amber force field ff19SB^26^ was used for protein and ion residues, Glycam06j-1^27^ for carbohydrates and TIP3P^28^ as a water model. Preliminary tests performed with other carbohydrate force fields showed that only Glycam06j correctly predicts the chair-to-boat transitions of the reactant states demonstrated by Petersen et al.^14^. In particular, using the CHARMM36 force field, the terminal Man persisted in a chair structure even with application of explicit restrains to facilitate a ring distortion. The input systems were energy-minimized using the steepest-descent algorithm with a tolerance of 1000 kJ mol^*−*1^ nm^*−*1^, restraining all heavy atoms. Consecutive equilibrations of 250 ps in an NVT ensemble with restrained heavy atoms, and a minimum of 1 ns in the NPT ensemble without restraints were performed prior to the production MD runs. The simulations were conducted using the leap-frog algorithm with an integration time step of 2 fs. The LINCS algorithm^29^ was applied to constrain bonds involving hydrogen atoms. Temperature control was achieved using the velocity-rescaling method^30^ with a time constant of 0.1 ps, maintaining a reference temperature of 310.15 K. The system pressure was set to 1 bar and controlled by the Parrinello-Rahman barostat with a time constant of 5 ps and a compressibility of 4.5*×*10^*−*5^ bar^*−*1^. For non-bonded interactions, the Verlet list scheme^31^ was used, updating the neighbor list every 100 steps. Electrostatic interactions were calculated using the Particle Mesh Ewald (PME) method^32^, employing a real-space cutoff distance of 1.2 nm.

### Enhanced sampling simulations of glycans (within CAZymes)

To ensure thorough phase-space sampling of glycans, especially within the catalytic site, enhanced-sampling simulations were carried out using a combination of the Hamiltonian replica exchange method with solute scaling (REST2)^33,34^ and the replica exchange with collective variable tempering (RECT) method^35^, which is based on well-tempered metadynamics (WT-MetaD)^36^. This combined approach, referred to as REST-RECT, was successfully developed specifically for glycans in a previous work^3^ and employed here for all simulated systems. Starting structures for REST-RECT were derived from equilibrium simulations, and the enhanced sampling runs conducted in a NPT ensemble, as previously described. The entire *N* -glycan was designated as the solute region, with its Hamiltonian scaled across *α* replicas using scaling factors *λ*_*α*_. These factors were applied to influence long-range electrostatics, Lennard-Jones interactions, and dihedral-angle interactions. Sixteen replicas were utilized, with *λ*_*α*_ values set to 1, 0.98, 0.95, 0.92, 0.90, 0.87, 0.84, 0.81, 0.78, 0.75, 0.72, 0.69, 0.65, 0.62, 0.58, and 0.55, corresponding to an effective temperature range from 310.15 K to 570 K. A geometric progression of *λ*_*α*_ values, as often employed in similar systems, proved less efficient for replica exchanges. Throughout the simulations, water and ions were maintained at the base temperature. The RECT component applied simultaneous biases to all dihedral angles through one-dimensional WT-MetaD bias potentials in each replica *α*. The dihedral angles were defined as *ϕ* = O5^*’*^–C1^*’*^–O*x*–C*x, ψ* = C1^*’*^–O*x*–C*x*–C(*x*–1) and *ω* = O6–C6–C5–O5, with *x* being the carbon number of the linkage at the non-reducing end. Replica-specific bias factors *γ*_*α*_ were defined as 1, 1.13, 1.27, 1.43, 1.61, 1.82, 2.05, 2.31, 2.60, 2.93, 3.30, 3.72, 4.19, 4.73, 5.32, and 6 along the replica ladder. Gaussian hills were deposited every *τ*_*G*_ = 1 ps, with a width of 0.35 rad and a height determined by *h*_*α*_ = (*k*_*B*_Δ*T*_*α*_*/τ*) *×τ*_*G*_, where *k*_*B*_ is the Boltzmann constant, Δ*T*_*α*_ = *T*_0_(*γ*_*α*_ *−* 1) is the boosting temperature, and *τ* = 4 ps is the bias evolution characteristic time. Replica exchanges were attempted every 400 steps and evaluated using the Metropolis-Hastings criterion.

Weak distance restraints (150 kJ/mol) were applied to keep the ions and glycan substrates located in the binding sites, particularly in higher-temperature replicas, preventing their diffusion. For MII, restraints included distances of 0.5, 0.2, 0.4, and 0.2 nm between the Zn^2+^ ion and amino acids H90 and H471, as well as atoms O2 and O3 of Man4. In simulations without the Zn^2+^ ion, restraints were not applied. For simulations at fixed D341-O6 distance, an harmonic restrain was applied between the H_*D*341_ and O6 atoms, with an equilibrium distance of 0.15 nm and a force constant of 5000 kJ mol^*−*1^ nm^*−*1^. For MI, restrained distances of 0.25 nm between the Ca^2+^ ion and T688, as well as between the Ca^2+^ ion and atoms O2 and O3 of Man7, were applied. Each replica was simulated for 500 ns, yielding a cumulative sampling time of 8 *µ*s. In contrast, enhanced sampling of free glycans in solution required only 12 replicas. For this case, a geometric progression of *λ*_*α*_ between 1 and 0.42 and *γ*_*α*_ between 1 and 14 was sufficient. Sufficient replica exchanges were validated by analyzing replica transitions along the ladder, calculating the probability of replica exchanges and round trip times. Additionally, convergence of glycan conformer distributions were assessed by checking the moving average of dominant glycan conformers over time, for consistency (see SI).

### Free energy reconstructions

We note that, apart from minor distance restrains, the ground replicas were unbiased due to their associated factors *λ*_0_ = 1 and *γ*_0_ = 1. Application of the weighted histogram analysis method (WHAM) in order to re-weight the ground-replica distributions under consideration of the influences from the distance restrains did not reveal any significant differences compared to directly using the unbiased ground replicas. We therefore neglected the usage of WHAM and used ground replicas directly in all subsequent analyses. Coordinates and variables were saved every 10 ps or more often, resulting in at least 50,000 data points per simulation. Free energies were calculated from the probabilities *P* according to the relation Δ*G* = *−k*_*B*_*T* ln(*P*). One-dimensional free energy profiles were calculated for the Cremer-Pople pucker coordinate *θ*, using block averaging in order to improve statistics, separating the data set in evenly distributed blocks (5 to 10 depending on the individual system). The average of all blocks 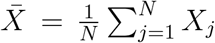 was calculated over *N* blocks, where *X*_*j*_ is the average calculated within each *j*th block. Error bars were calculated as standard deviations of the sampling distributions (standard error of the mean):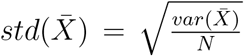, with the variance of the sampling distributions 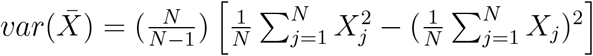. Twodimensional pucker histograms along *ϕ* and *θ* with 200 *×* 200 bins were constructed using PLUMED and plotted using the Mollweide (also termed elliptical) projection. This pseudocylindrical map projection fullfils the equal-area condition, meaning that areas, densities and, thus, free energy values, are preserved in the 2D representation. The reduced conformational phase space of a glycan was represented in a latent space by principal component analysis (PCA), calculated using the scikit-learn package. ^37^ All sin and cos values of dihedral angles of each glycan were defined as features (*n*_*features*_), resulting in a feature matrix **X** with shape *n*_*samples*_ *× n*_*features*_. This correctly accounts for the dihedral angle periodicity. Whenever two **X** from separate simulations were compared to each other, e.g. unbound versus bound glycan, the corresponding data sets were concatenated prior to PCA calculation. Free energies (Δ*G*) along the principal components 1 and 2 defining the low-dimensional latent-space matrix **T** were calculated by constructing two-dimensional histograms with 35 bins and converting the histogram probabilities *P* according to the free energy equation above.

### Steered and restrained MD simulations

Steered MD simulations were performed for native MII with bound M5G0, using a snapshot from the REST-RECT simulations as starting configuration where M5G0 adopts the #1* conformer (*vide infra* for the conformer labelling). A minimization and subsequent production run of 6 ns simulation in the NPT ensemble were performed with the same simulation parameters as before. The PLUMED package with its module MOVINGRESTRAINT was employed to apply harmonic restraints to certain variables in order to artificially alter the conformation of the glycan. Two different sets of steered MD simulations were performed. In the first set, restraints were applied to *ϕ, ψ* and *ω* in the Man2-Man3 linkage as well as to *ψ* and *ω* in the Man3-Man4 linkage, with force constants of 500 kJ mol^*−*1^ nm^*−*1^ for all angles. Especially these dihedral angles have been chosen because they clearly distinguish between the #1* and #2* conformer. In Man2-Man3, *ϕ* was varied between 1.0 and 3.0 rad, *ψ* between 3.0 and -2.0 rad, *ω* between 3.0 and 1.0 rad. In Man3-Man4m, *ψ* was varied between 1.0 and 3.0 rad and *ω* between 1.0 and -1.0 rad. In the second set, solely *θ* of Man4 was varied between 0.25 and 1.5 rad to drag the system from the ^4^*C*_1_ chair to a distorted boat/skew-boat. Each steered-MD cycle lasted 3 ns, with 0.5 ns of equilibration, followed by 0.5 ns of ramping and 1 ns of equilibration, prior to 0.5 ns of ramping back to the original conditions and 0.5 ns of final equilibration. Variables were outputted and plotted every 10 ps.

The simulation of M5G0 restrained in its #2* conformer was enabled by the application of harmonic restrains on all dihedral angles using force constants of 50 kJ mol^*−*1^ nm^*−*1^. Sampling of puckering landscapes was achieved using REST2, without any explicit biases on dihedral angles, as these were fixed. The same number of replicas and scaling parameters were used as already described for free glycans in solution.

### Correlation analysis

The Pearson correlation coefficients were computed using the pairwise correlation method implemented in Pandas. Pairwise correlations between dihedral angles and pucker coordinates were calculated and combined in a correlation matrix. As for the PCA analysis, dihedral angles were converted into their sin and cos values, in order to account for angle periodicity. For the correlations between glycan conformers and pucker coordinates, first the conformers were defined based on the glycan dihedral angles and individual labels were assigned by means of the GlyCONFORMER package.^3^ As conformer labels are ordinal, meaning categorical without a known distance between them, one-hot encoding was employed to convert conformer values into binary format and subsequently compute pairwise Pearson coefficients.

### Classifying glycan conformations

The GlyCONFORMER code identifies and classifies glycan conformers from MD simulations by assigning a unique string based on adopted values of glycan dihedral angles.^3^ Each glycosidic linkage contributes at least two dihedral angles (*ϕ* and *ψ*), with an additional *ω* angle for 1–6 and 2–6 linkages. A conformer is represented by a digit string of length *n* equal to the number of dihedral angles in the glycan. Hereby, each digit corresponds to a dihedral angle interval, following the IUPAC nomenclature.^38^ The string starts at the free reducing end and includes digits for *ϕ, ψ*, and *ω* (if present), in this order. At branching junctions (e.g., *α*1–6 and *α*1–3), a separator labeled by the C atom at the branch origin (e.g., **6**–) is added. The string proceeds along the higher C atom branch (e.g. 6) to the terminal residue before returning to follow the lower branch (e.g. 3). The assignment involves calculations of the free-energy profiles along each dihedral angle from converged (typically, enhanced-sampling) MD trajectories. The local minima in these profile are labeled according to the nomenclature above, and the same label is inherited by all angles in each free-energy basin around the minimum.

## Results and Discussion

### The reactant state: how is it reached?

We start our investigation by following the pucker itinerary of the terminal Man4 of M5G0, at subsite *-1* with respect to the glycosidic bond cleavage, when inserted in the catalytic site of MII from *D. Melanogaster* (PDB: 3CZN, see Methods). The conformational phase space of M5G0 calculated with the REST-RECT methodology^3^ (Figure S1 S8) reveal that Man4 adopts a ^4^*C*_1_ chair when the glycan is free in solution (Figure 3 A). Upon binding to MII, the puckering landscape of Man4 is altered, acquiring a second minimum around the *E*_5_/^0^*H*_5_ envelope/half-boat (Figure 3 B). The minimum corresponds to the ring shape that was experimentally determined for the acid-base MII mutant D341N^15^ and is close to the reactant-state ^*O*^*S*_2_/*B*_2,5_ described by Petersen et al.^14^. Binding to MII leads to a 30 kJ/mol reduction in energy barrier for a ring-distortion along *θ* from ^4^*C*_1_ towards the equatorial region, which is now only about 7 kJ/mol higher in energy (Figure 3 A/B). The experimentally observed hydrogen bond O_*D*92_-H_*O*2_ is properly formed (Figure S9 A).^14^ However, the D341-O6 distance between the center of mass of the carboxylic group of the catalytic acid/base and the oxygen atom in the glycosidic linkage is about 0.4 nm, which is larger than what is necessary to fully distort the ring.^14^ Possibly, the fixed-charged force field employed in our simulations underestimate this distance because of the lack of polarizability of the residues. We therefore apply a distance restrain on H_*D*341_-O6 (Figure S9 B) to keep it at 0.15 nm, which leads to a broadening and shift of the global energy minimum towards the reactant state ^*O*^*S*_2_/*B*_2,5_ (Figure 3 C).

**Figure 3.**
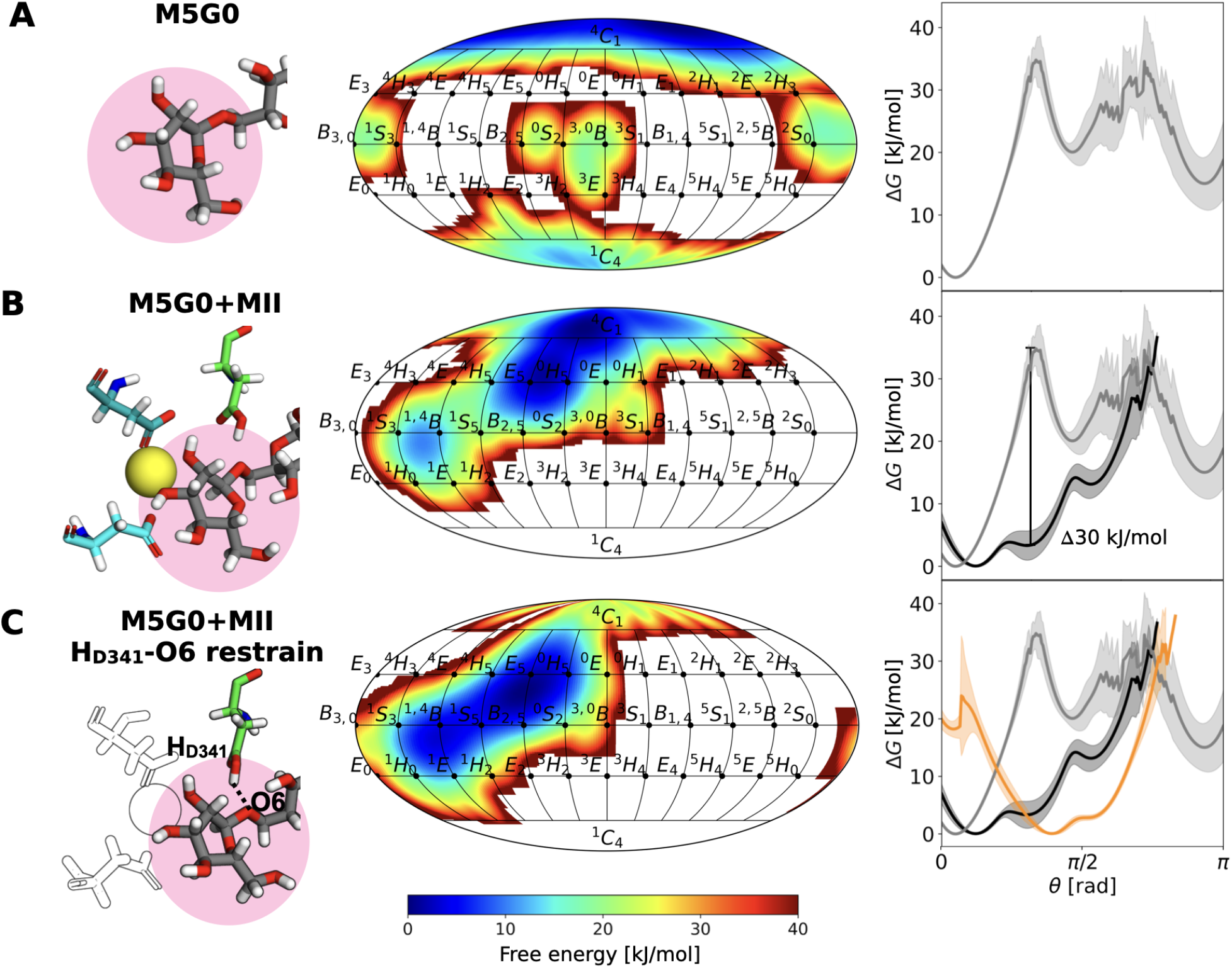
Ring puckering in different chemical environments. Ring distortion of the terminal Man4 at subsite *-1* (highlighted in pink in the atomistic snapshot) in M5G0 monitored by the 2D Cremer-Pople representation along *ϕ* and *θ* as well as 1D along *θ* in different chemical environments: **A** M5G0 in aqueous solution (gray), **B** M5G0 in the catalytic site of MII (black), **C** M5G0 in the catalytic site of MII with a distance restraint (dotted black line in the atomistic snapshot with only important residues highlighted by colors) between O6 of M5G0 and the H atom of the carboxyl group of D341 (orange). Error bars are plotted as shaded regions, representing the standard deviation calculated from block averaging.

These simulations show that the binding of M5G0 into the active sites activates Man4 into a distorted shape prone to chemical attack. However, in contrast to earlier claims that the *B*_2,5_ structure is necessary for the mannose residue to fit into the binding pocket,^15^ our simulations reveal that ^4^*C*_1_ does also perfectly fit. The ring distortion is therefore not a prerequisite for, but an effect of the confinement within the catalytic site.

### The reactant state: how is it influenced?

The specificity of protein-ligand interactions responsible for the ring distortion was investigated by studying the effect of various amino-acids mutations and deletion of the Zn^2+^ ion on the ring puckering. ^15^ REST-RECT simulations were performed for four mutants: D341A, D204A, D92A and D472A. These amino acids were selected based on the variation of their distances from Man4 depending on the ring shape (^4^*C*_1_ or *E*_5_/^0^*H*_5_), as observed in unbiased MD simulations (Figure S10). The D341A mutation removes the catalytic acid/base residue from the active site, experimentally leading to an almost inactive MII enzyme (actually observed for D341N, PDB: 3BUP), although a distorted mannose at subsite *-1* persists. ^15^ Our simulations confirm these findings, leading to only a slight energy increase of the intermediate state *E*_5_/^0^*H*_5_ compared to the wild-type MII simulation (Figure 4 A). It becomes apparent that D341 is primarily crucial as a hydrogen donor for subsequent covalent complex formation with Man4, but not necessary to activate the ring distortion.

**Figure 4.**
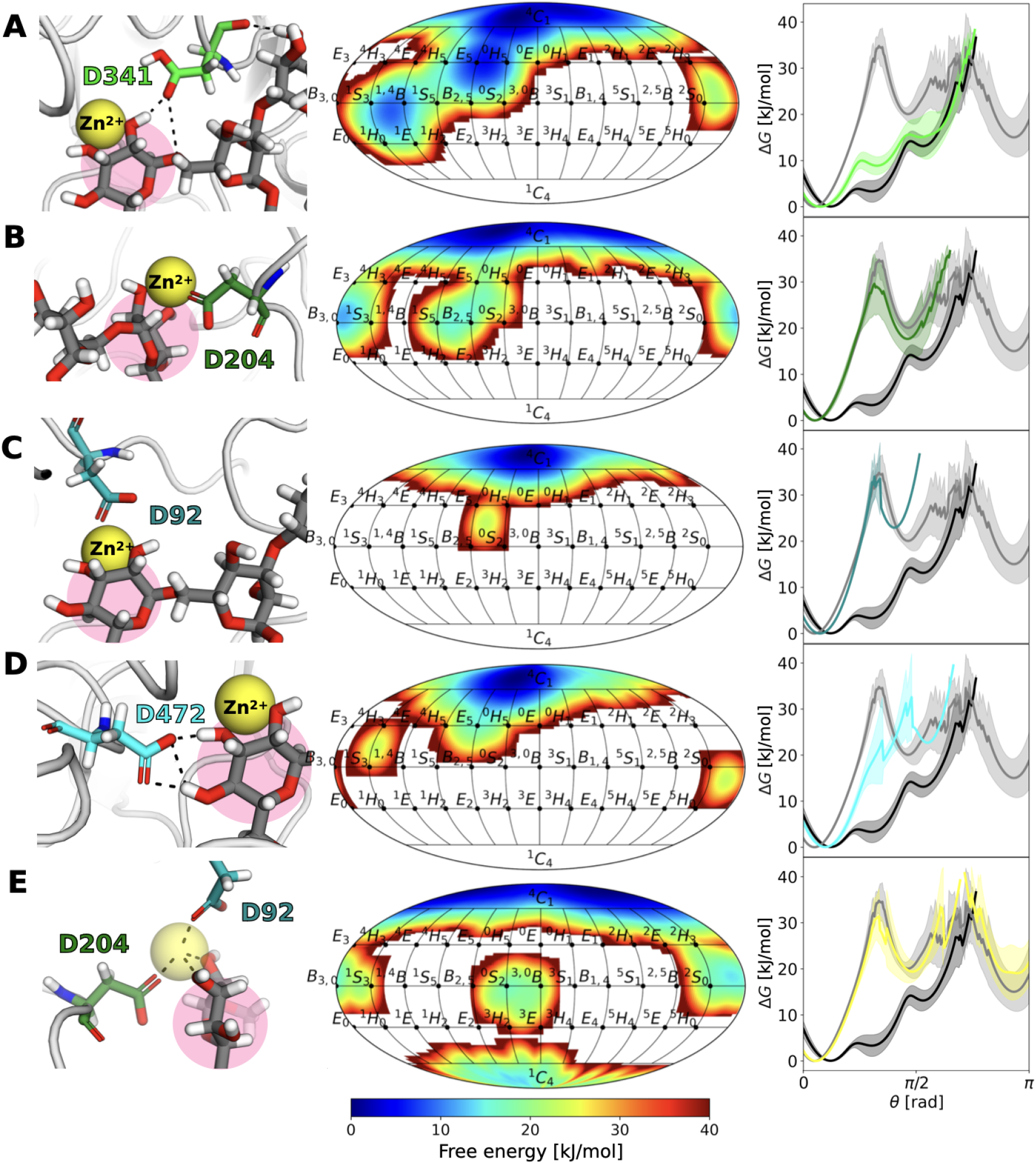
Mutants altering ring distortion. For **A** D341A, **B** D204A, **C** D92A, **D** D472A, **E** absence of the Zn^2+^ ion, the following separate panels are shown: atomistic snapshot of the catalytic site under native conditions (left), free energy surface along the Cremer-Pople puckering parameters *θ* and *ϕ* for the terminal Man4 of glycan M5G0 (highlighted in pink in the atomistic snapshot) bound to mutant MII (center), and free energy profile along *θ* of Man4 for the unbound (gray), bound (black), and altered state (colored) (right).

Mutation of the catalytic nucleophile D204A experimentally led to the same catalytic inactivation as D341N, however without Man4 being distorted away from its solution structure ^4^*C*_1_ (PDB: 3CZN). In our simulation, the free energy profile along *θ* resembles that of a free glycan in solution, loosing the local minimum for the intermediate states and therefore being in agreement with experiments (Figure 4 B). Mutations D92A and D472A, that have not been tested experimentally so far, have the effect of loosing either the correct positioning of the Zn^2+^ ion or hydrogen bonds with the Man4, which thus presented only a global ^4^*C*_1_ minimum (Figure 4 C/D). These latter mutants completely compromise the ring distortion, and are thus likely to be catalytically inactive.

The role of the Zn^2+^ ion in the catalytic activity of MII has long been debated, putatively helping in activating the ring distortion or stabilizing the electronic charge shifts crucial for reaching the reaction’s transition state. Interestingly, our simulations show that removal of the Zn^2+^ ion prevents ring distortion completely, leading to an energy profile identical to that of free M5G0 in solution (Figure 4 E). This is in line with experiments showing no activity for MII lacking Zn^2+^ in the catalytic site and confirms its direct role in ring distortion. ^39^ However, since electronic polarization is not implemented in our force-field, the action is not due to a charge redistribution within the ring.

### The reactant state: how do glycan conformations matter?

The simulations presented in the last section show that the smallest change in the interior of the catalytic site has a large effect on the activation of the glycan substrate responsible for bringing about a conformationally reactive state. The term conformation in the context of CAZyme’s catalytic mechanisms predominantly refers to ring puckering, whereas the overall glycan shape has been largely overlooked, despite the importance of ligand flexibility in protein-ligand interactions and docking affinity. Using our enhanced-sampling protocol, we can now focus on the distributions of sugar conformers associated with changes in the dihedral angles along the glycosidic bonds. PCA-reconstructed two-dimensional free energy surfaces that represent the accessible conformational phase space reveal that M5G0 in solution can adopt multiple conformers (#1-#6) that differ primarily in the *ω* angles of the 1-6 linkages (Figure 5 A and S1). In stark contrast, M5G0 bound to MII is confined to just two dominant global conformers, #1* and #2* (Figure S2), which differ significantly from the conformers observed in solution (Figure S11). This shift reflects the tight binding of almost all monosaccharide residues in M5G0 to the catalytic, holding, and anchor site of MII.^40^

**Figure 5.**
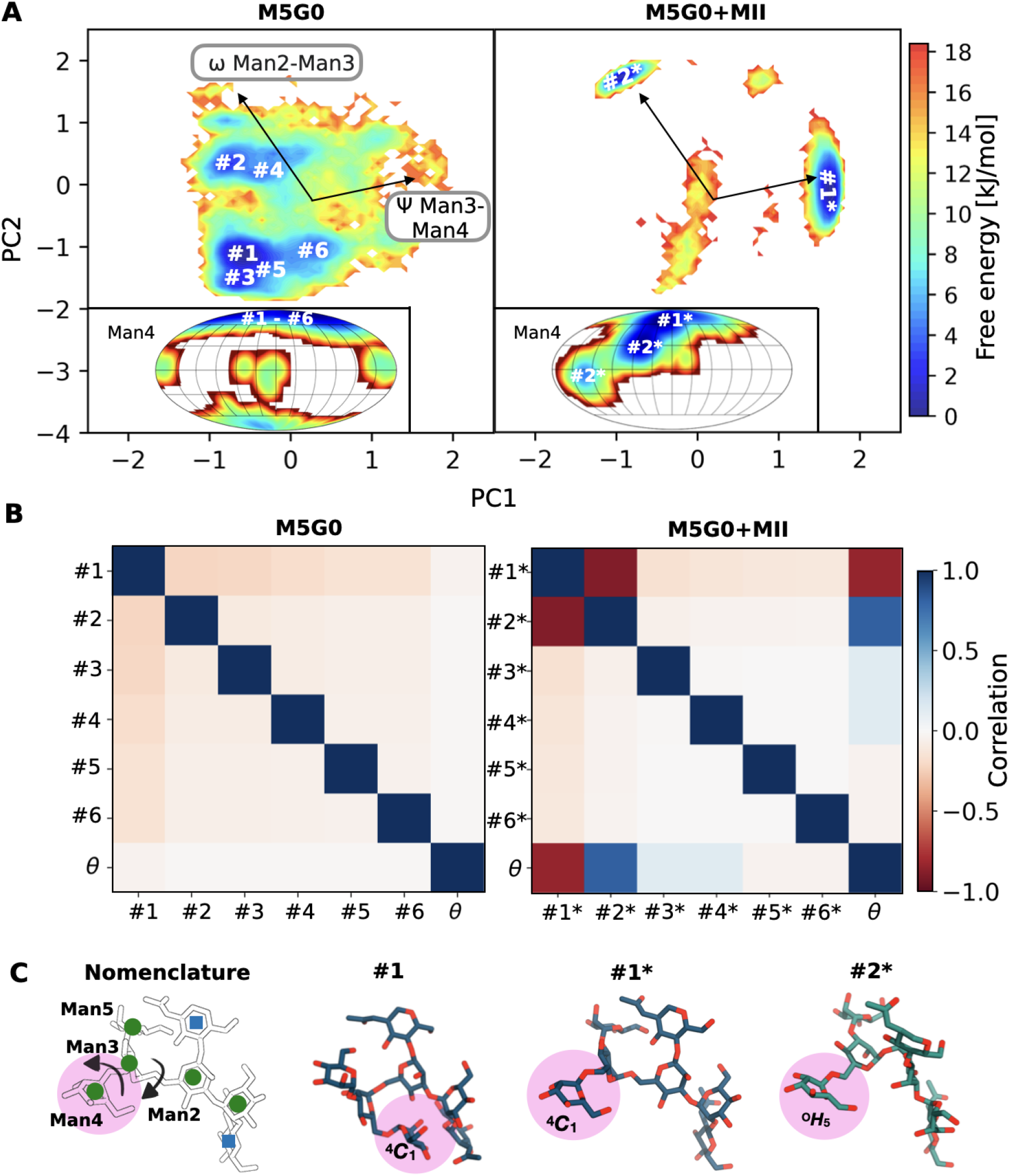
Dihedral angle-puckering correlation of M5G0 in MII. **A** Comparative free energy surfaces of the conformational phase space for M5G0 in solution and bound to MII projected along PC1 and PC2. Vectors indicate the original feature axes with highest variance, pointing in the direction with highest squared multiple correlation with the principal components. The inserts show the corresponding free energy surfaces along *θ* and *ϕ* for the terminal Man4, with minima labeled by the respective global glycan conformers adopted in that puckering region. **B** Correlation matrix for glycan conformers and *θ* of Man4 for M5G0 in solution and bound to MII, utilizing the Pearson correlation coefficient. **C** Atomistic snapshot of M5G0, highlighting the differences between conformer #1, #1* and #2*, as indicated by arrows in the schematic structure on the left. Conformer details in Figure S1 - S2.

We identified a striking correlation between this shift in global conformation and the ring distortions of Man4. In solution, all #1-#6 M5G0 conformers present a ^4^*C*_1_ ring shape for Man4 (Figure 5 A inserts). After binding, the ^4^*C*_1_ chair is preserved only for the #1* conformer, whereas in the #2* conformer, Man4 experiences a transition to the half-boat ^0^*H*_5_. This correlation was quantified by computation of the pairwise Pearson correlation coefficients between global glycan conformers (defined by their dihedral angle values) and the pucker coordinate *θ* of Man4 (Figure 5 B). The dihedral angle and puckering degrees of freedom are all completely uncorrelated for M5G0 in solution. In contrast, MII-bound M5G0 exhibits strong positive and negative correlations between *θ* and conformers #2* and #1*, respectively.

Further detailed analyses of correlations between dihedral angles and ring shapes (Figure S12) revealed specific dihedral angles, primarily within the Man2-Man3 and Man3-Man4 linkages, that strongly correlate with *θ* of Man4 and with each other in MII-bound M5G0 (Figure 5 C). Structural comparisons of the conformers clearly show that the shift from #1 to #1* is driven by changes in the Man3-Man4 dihedral angle, leading to a complete rearrangement of the entire glycan branch (Figure 5 C). Similarly, the transition from #1* to #2* involves a structural reorganization of the Man2-Man3 glycan branch, repositioning it relatively to the remaining core structure. This shift coincides with the adoption of the half-boat ^0^*H*_5_.

Analysis of M5G0 conformers in mutated MII variants reveal a distinct preference for the #1* conformer, accompanied by a significant reduction in the #2* conformer. This observation aligns with a decrease in ring distortion which is known for at least some mutants to result in a loss of catalytic activity^15,40^, emphasizing the correlation between linkage and ring distortions (Figure S13). We further restricted the 3D structure of a free M5G0 glycan in solution to the #2* conformation, in order to test if solely the dihedral angles defining the glycan conformation may be responsible for the shift in pucker propensity. However, in this case the Man4 remains in the ^4^*C*_1_ chair, suggesting that binding to the catalytic site is necessary to both restricting the glycan conformers and activating the ring distortion (Figure S14).

To better investigate the causality underlying the formation of the reactant state, we conducted two sets of steered MD simulations: a first one by alternating between conformers #1* and #2* using restraints on selected dihedral angles of M5G0 (Figure 6 A), and a second one by varying *θ* values of Man4 between a ^4^*C*_1_ chair and boat/skew-boat ring structure (Figure 6 B). Particular attention was given to the D341-O6 distance, as shorter distances were observed to promote a full chair-to-boat distortion of Man4 (Figure 3C). Our results reveal that, upon inducing a transition to the #2* conformer, Man4 concurrently shifts to the half-chair ^0^*H*_5_. This transition is accompanied by a reduction in the D341-O6 distance from 0.5 nm to 0.4 nm (Figure 6 C. Forcing a Man4 ring distortion as a constraint had a much smaller effect (Figure 6 B). Specifically, only a change of the Man3-Man4 *ψ* dihedral, but no change of the *ω* dihedral was observed. Additionally, the D341-O6 distance experienced barely visible variations. We can thus conclude that the activation of Man4 is the consequence of the following events. First, binding into the catalytic site of MII imposes a strong constraint on the dihedral-angle conformational phase space of M5G0. This dihedral shift then drives a ring distortion at subsite *-1*, which leads to a reduced D341-O6 distance, as required by the acid-base reaction initiated by a proton transfer from D341 to O6. Importantly, ring distortion not only facilitates the development of the OCI character of the *-1* sugar, but also ensures precise positioning of the glycosidic linkage to be cleaved within the catalytic site (Figure 6 C).

**Figure 6.**
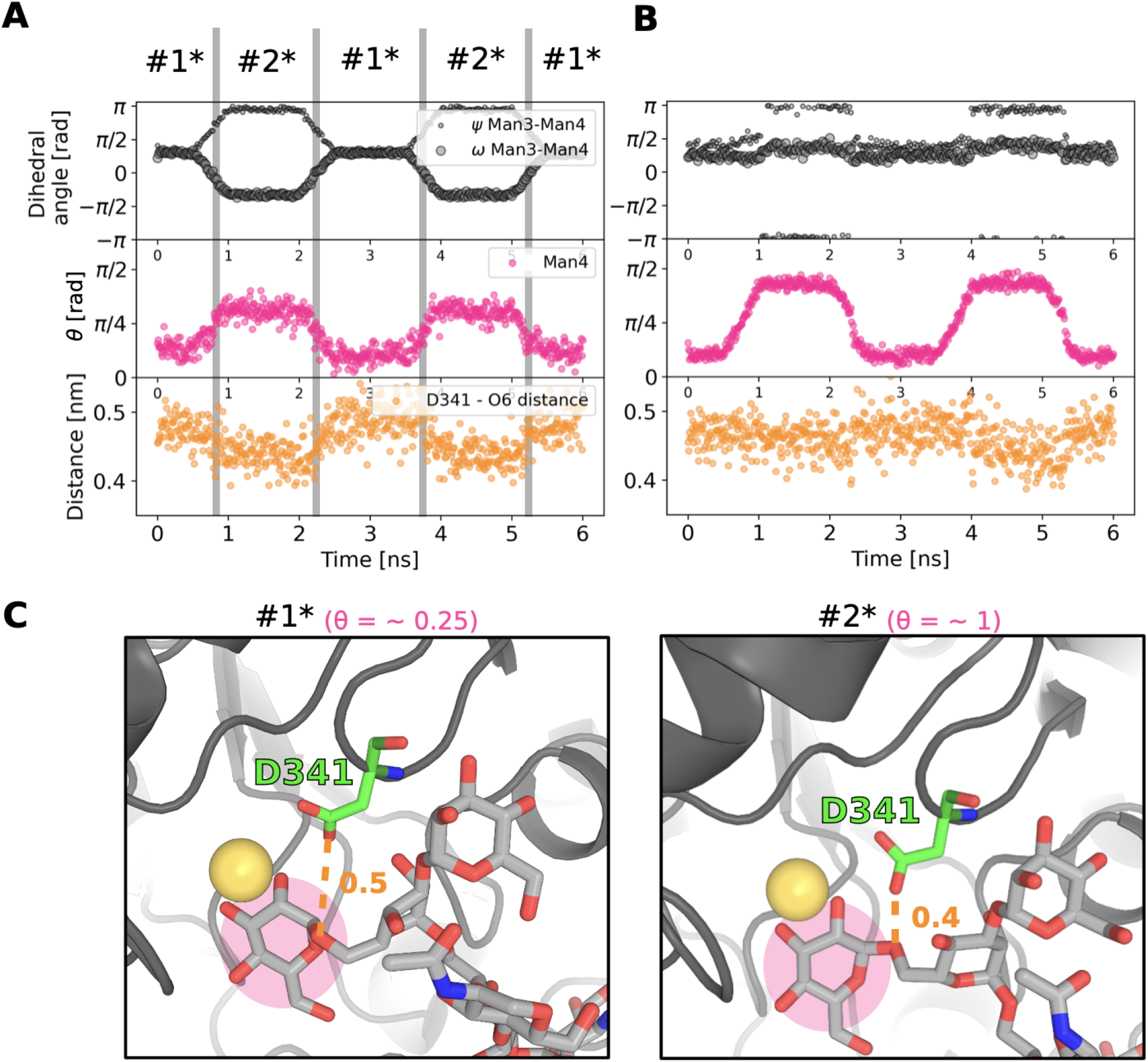
Local conformer #2* induces ring distortion and shortens D341-O6 bond. Monitoring of selected dihedral angles, ring distortion of Man4 (*θ*) and D341-O6 distance over steered MD simulations of native MII where **A** dihedrals of M5G0 are altered to switch between conformers #1* / #2* and **B** *θ* of Man4 is altered between chair ^4^*C*_1_ and boat/skew-boat structures. **C** Atomistic snapshots of **A** in the #1* and #2* conformers, respectively, highlighting the difference in D341-O6 distance associated with reorientation of dihedral angles and therefore conformers. M5G0 is shown in licorice with gray carbon atoms, D341 in green, the Zn^2+^ ion as a yellow sphere and the protein in cartoon style in dark gray. The subsite *-1* is highlighted by a pink circle.

### The reactant state: The reason behind substrate specificity

The knowledge we have gained about the factors governing the formation of the reactant state helps us explain other aspects of the MII catalytic mechanism. In particular, a comprehensive understanding of the substrate binding selectivity is urgently needed in the context of cancer treatment to design effective inhibitors for MII which do not bind to the similar lysosomal *α*-mannosidase (L-MII). Previous studies have highlighted that a specific MII region, characterized by amino acids Q64, Y267, H273, P298, W299, and R410 (Figure 7 A), plays a crucial role in the binding selectivity of MII toward M5G0. Specifically, the terminal GlcNAc7 residue, which differentiates M5G0 from other high-mannose-type N-glycans, binds to this region, thus referred to as the anchor site.^40^ Experimental evidence has shown that the smaller M5 substrate, lacking the terminal GlcNAc7 residue, can bind to MII in the correct position. However, the glycolytic activity of MII towards M5 is reduced by a factor of 80 compared to M5G0 for yet unknown reasons. ^15,40^ To reveal the relationship between anchor site binding and catalytic efficiency, we modeled the conformational phase space of M5 before and after binding to MII (Figure 7). ^40^ REST-RECT simulations of M5 in solution revealed a predominant ^4^*C*_1_ chair for Man4, whereas M5 bound to MII exhibited a ring distortion towards ^0^*H*_5_, similar to the distortion experienced in M5G0 (Figure 7 C/D). This indicates that the GlcNAc7 residue in the anchor site is not essential for inducing a ring distortion at subsite *-1*. Yet, M5 bound to MII displays a significantly broader conformational phase space, with five conformers exceeding a 5% probability (labelled #1^$^ to #5^$^). This is similar to the amount of conformers populated by M5 in solution, although different in shape (Figure S15 and S16). Notably, the #4^$^ conformer in M5+MII corresponds to the #1* conformer of M5G0+MII, and #3^$^ corresponds to #2* (Figure 7 E). Consistently, #4^$^ exhibits a negative correlation to the *θ* puckering coordinate of Man4 (Figure 7 F), as also observed for #1*. However, none of the other conformers correlate with *θ* of Man4, suggesting that ring distortion is excluded for #4^$^, but possible for any other conformer independently of their dihedral conformations. Notwithstanding this possibility, the most populated conformers of M5 differ in the conformation of the important glycosidic bond Man2-Man3, potentially hindering the efficient rearrangement of the linkage for cleavage.

**Figure 7.**
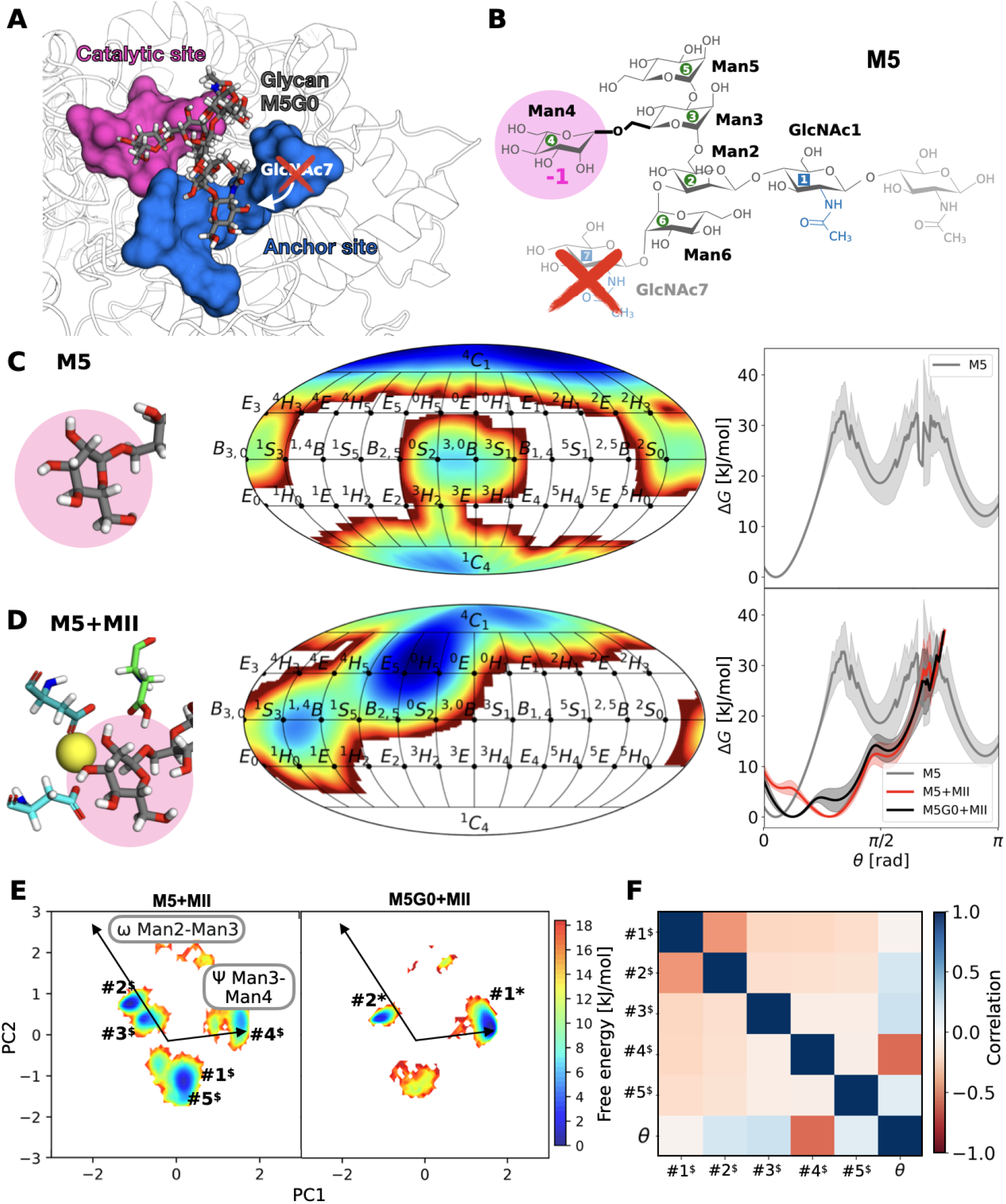
Unoccupied anchor site results in larger conformational phase space for M5 bound to MII. **A** Atomistic snapshot of catalytic (pink) and anchor site (blue) in MII with bound M5G0. Removal of GlcNAc7 results in M5 and an unoccupied anchor site. **B** Atomistic structure and nomenclature of M5 with the lacking GlcNAc7 shown in transparent and marked by a red cross. **C** Ring distortion of the terminal Man4 in glycan M5 monitored in a 2D representation along *ϕ* and *θ* as well as 1D along *θ* in aqueous solution (gray). **D** Same, for M5 bound to MII (red), compared to the M5G0+MII system (black). **E** Comparative free energy surfaces of the conformational phase space for M5+MII and M5G0+MII projected along PC1 and PC2. **F** Pearson correlation matrix for M5 conformers and *θ* of Man4 bound to MII.

These findings underscore the anchor site’s critical role in reducing the substrate flexibility in the binding site, stabilizing correlated glycan conformers and ring puckering that efficiently promote the glycosidic bond cleavage. Interestingly, L-MII lacks such an anchor site, opening up a novel inhibitor design strategy. Namely, MII-selective inhibitors could be designed to strongly bind to the anchor site, while simultaneously mimicking the #2* conformation of M5G0 in the catalytic site. This dual functionality would ensure tight binding and selectivity towards MII, thus reducing the probability of binding to L-MII.^40^

#### Extension to other CAZymes

The methodology and workflow that we have developed can be seamlessly applied to other CAZymes than MII, enabling a deeper understanding of their own specificity and catalytic mechanisms. As an example, we investigated the related MI enzyme, which cleaves the high-mannose type N-glycan M9 (Figure 8 B), utilizing a Ca^2+^ ion, E599 as the catalytic base, and E330 as the catalytic acid within its active site (Figure 8 A). ^41^ This enzyme is critical for the maturation of N-linked oligosaccharides in mammalian cells and participates in the degradation of misfolded glycoproteins.^41–44^ Unlike MII, which targets the flexible *α*1-6 linkage, MI specifically hydrolyzes the *α*1-2 glycosidic bond between Man6 and the terminal monosaccharide Man7 at subsite *-1* (Figure 8 B).^45^ REST-RECT simulations of M9 before and after binding reveal critical differences in the ring distortion behavior of Man7. In solution, the global ^4^*C*_1_ chair is dominant (Figure 8 C). Binding to MI induces a shift to the ^1^*C*_4_ chair (Figure 8 D), consistent with experimental observations ^45^, and increases significantly the propensity for the reactant-state boat ^3,0^*B*.^46^ Upon binding to MI, the conformational phase space of M9 is notably restricted to one dominant conformer (#1*) with minimal contributions from other conformers (Figure S17 S19). Importantly, the glycosidic bond between Man6 and Man7, formed via the O3 atom, remains in the same conformation regardless of ring distortion and conformer (Figure 8 E). This is further supported by the absence of correlation between ring distortion in Man7 and glycosidic bond orientation for all conformers (Figure 8 F). Our finding underscores a distinct catalytic mechanism that does not rely on correlations between dihedral angles and puckering degrees of freedom in the case where structural rearrangements of the bond to be cleaved are unnecessary.

**Figure 8.**
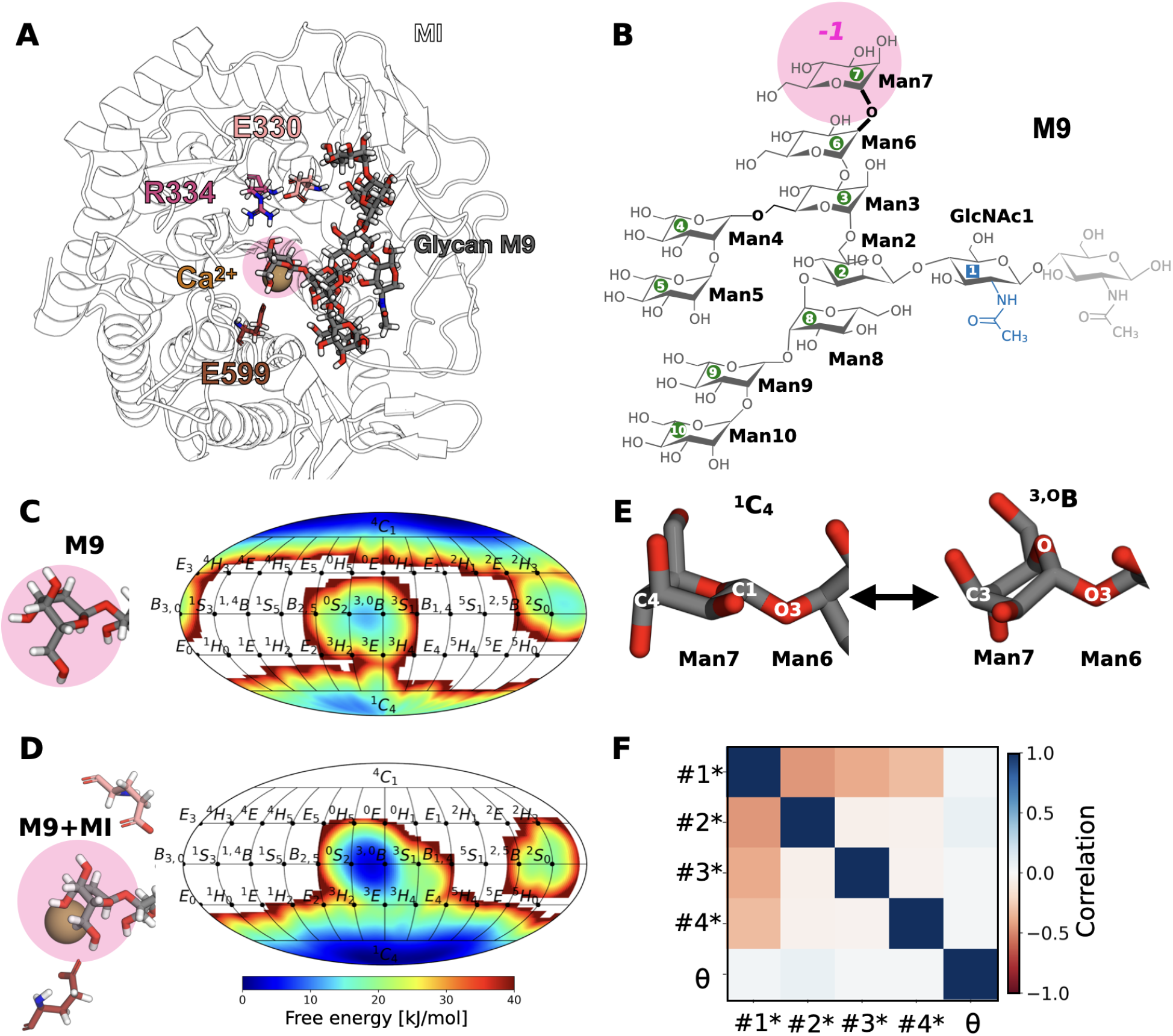
No correlation between ring distortion and glycan conformers for M9 in MI. **A** Atomistic structure of MI with bound glycan M9, Ca^2+^ ion and catalytic residues E330, R334 and E599. **B** Atomistic structure and nomenclature of M9 with labeling of linkages according to neighboring residue numbers. The Man6-Man7 linkage is highlighted in bold and is to be cleaved by MI. **C** Ring distortion of the terminal Man7 at subsite *-1* in glycan M9 monitored in a 2D representation along *ϕ* and *θ* in aqueous solution. **D** Same, for M9 in the catalytic site of MI. **E** Details of the orientation of the Man6-Man7 glycosidic bond in the two main puckering states. **F** Correlation matrix for M9 conformers and *θ* of Man7 bound to MI.

**Figure 9.**
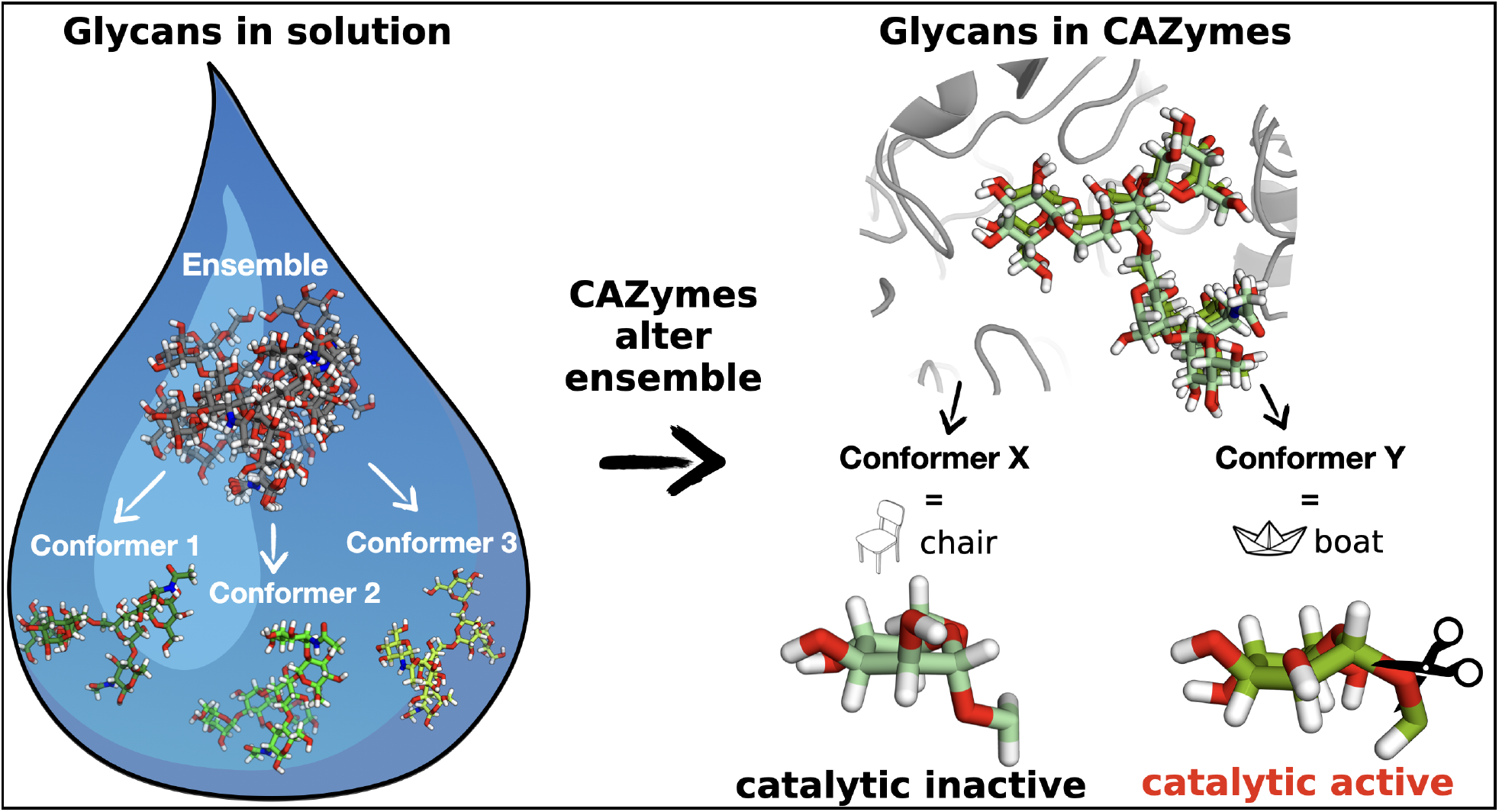
For Table of Contents Only.

## Conclusions

It is well-established that CAZymes from different families with similar substrate specificities employ diverse catalytic strategies. ^5,9^ In this study, utilizing enhanced-sampling molecular dynamics simulations with the REST-RECT methodology, we have provided detailed molecular insights into the diverse pathways that lead to stabilisation of glycan reactant states in mannosidases. Comparing three different substrates and two different enzymes, we demonstrated that the essential chair-to-boat ring distortion at subsite *−*1 is not achieved by a universal mechanism, but via individualized cascades of events.^45^

In all cases, the conformational phase-space of the substrates is significantly restricted to a few conformers upon entering the catalytic site. The conformer distribution is very system-dependent and is altered by even small modifications of the substrate, such as the absence of one terminal saccharide in M5 with respect to M5G0, or of the enzyme, such as mutation of single amino acids in the catalytic site. Specifically, MI locks its substrate M9 in only one major conformer, whereas M5G0 can adopt two distinct conformers in the binding pocket of MII, which differ substantially from those adopted in solution.

Upon binding, the terminal monosaccharide at position *-1* with respect to the bond-cleavage site increases its propensity for distorted ring puckering in all studied substrate/enzyme systems. In M5G0 bound to MII, the ring distortion is strictly correlated with the adoption of a certain global glycan conformation. This reveals a wholly new relationship between dihedral angles and puckering degrees of freedom that promotes an orientation of the glycosidic bond to hydrolytic cleavage. This correlation is observed for only one of the possible conformers of M5 bound to MII, and not observed for M9 bound to MI. In the latter case, the correlation is also not necessary, given that only one conformer is locked into the binding pocket with an already favorable orientation of the to-be-cleaved glycosidic bond. For M5G0 in MII, the correlation is crucial, because only in the distorted ring conformation the D341O6 distance is short enough to promote the initial proton transfer and kick-off the glycolytic reaction.

Rather than just providing a convenient environment for a static binding pose, the specific amino acid composition of the catalytic site and the pivotal presence of a divalent cation (Zn^2+^ in MII, Ca^2+^ in MI) influence the conformational dynamics of the substrates, opening up convenient channels towards the reactant states with oxocarbenium ion-like character of the sugars at subsite *-1*.^9^ Our findings emphasize the need for accessing the full conformational space, rather than determining only static snapshots, when attempting to fully understand the specificity and catalytic efficiency of CAZyme/substrate pairs. As different CAZyme families feature different mechanisms, tailored studies are needed that take into account the unique structural and chemical features of individual enzymes and substrates.

The methodologies and nuanced understanding presented in this study provide a valuable framework for the rational advancing of drug development strategies aimed at modulating CAZyme activities. For instance, by designing relatively large drug molecules that simultaneously bind both to the anchor site and the catalyitc site of MII with the specific #2* conformation, an efficient inhibition of MII could be achieved while minimizing a concomitant unwanted binding to L-MII.^8^ By aiming for precise ligand conformations that resemble certain steps in the catalytic itinerary of substrates, future CAZyme inhibitor designs could greatly enhance therapeutic applications, particularly in cancer treatment and other glycosylation-related diseases.

## Associated content

The generation of glycan conformer strings and conformer distribution plots was done by GlyCONFORMER, a python-based package deposited on GitHub under https://github.com/IsabellGrothaus/GlyCONFORMER. PLUMED input files for REST-RECT and steered MD simulations were deposited in the PLUMED-NEST repository under PlumID:25.007. Structure and trajectory files for all simulations can be accessed from https://doi.org/10.5281/zenodo.14858832. For REST-RECT simulations, only the ground replica trajectories are provided.

## Acknowledgement

The authors gratefully acknowledge the HPC resources provided by the National High Performance Computing Center Zuse-Institute Berlin (NHR@ZIB) under project ID: hbb00004. Additionally, the authors gratefully acknowledge the scientific support and HPC resources provided by the Erlangen National High Performance Computing Center (NHR@FAU) of the Friedrich-Alexander-Universität Erlangen-Nürnberg (FAU) under the NHR project ID: 22459. NHR funding is provided by federal and Bavarian state authorities. NHR@FAU hardware is partially funded by the German Research Foundation (DFG) – 440719683.

## Supporting Information Available

The Supporting Information file contains the GlyCONFORMER analysis for glycans simulated either in solution or bound to one of the CAZymes, including the glycan conformer distribution, convergence of glycan conformers during REST-RECT simulations and latent space projection via PCA of the conformer distribution. Furthermore, it provides additional plots for important protein-glycan distances and 2D free energy profiles of conformer distributions for several simulation systems. Detailed correlation matrices of dihedral angles and pucker coordinates are provided for M5G0 and M5G0+MII.

## Footnotes

1 Carbohydrate Active Enzymes database accessed 9th December 2024, http://www.cazy.org

