## Supporting Information for "Shaping the Glycan Landscape: Hidden relationships between linkage and ring distortions induced by carbohydrate-active enzymes"

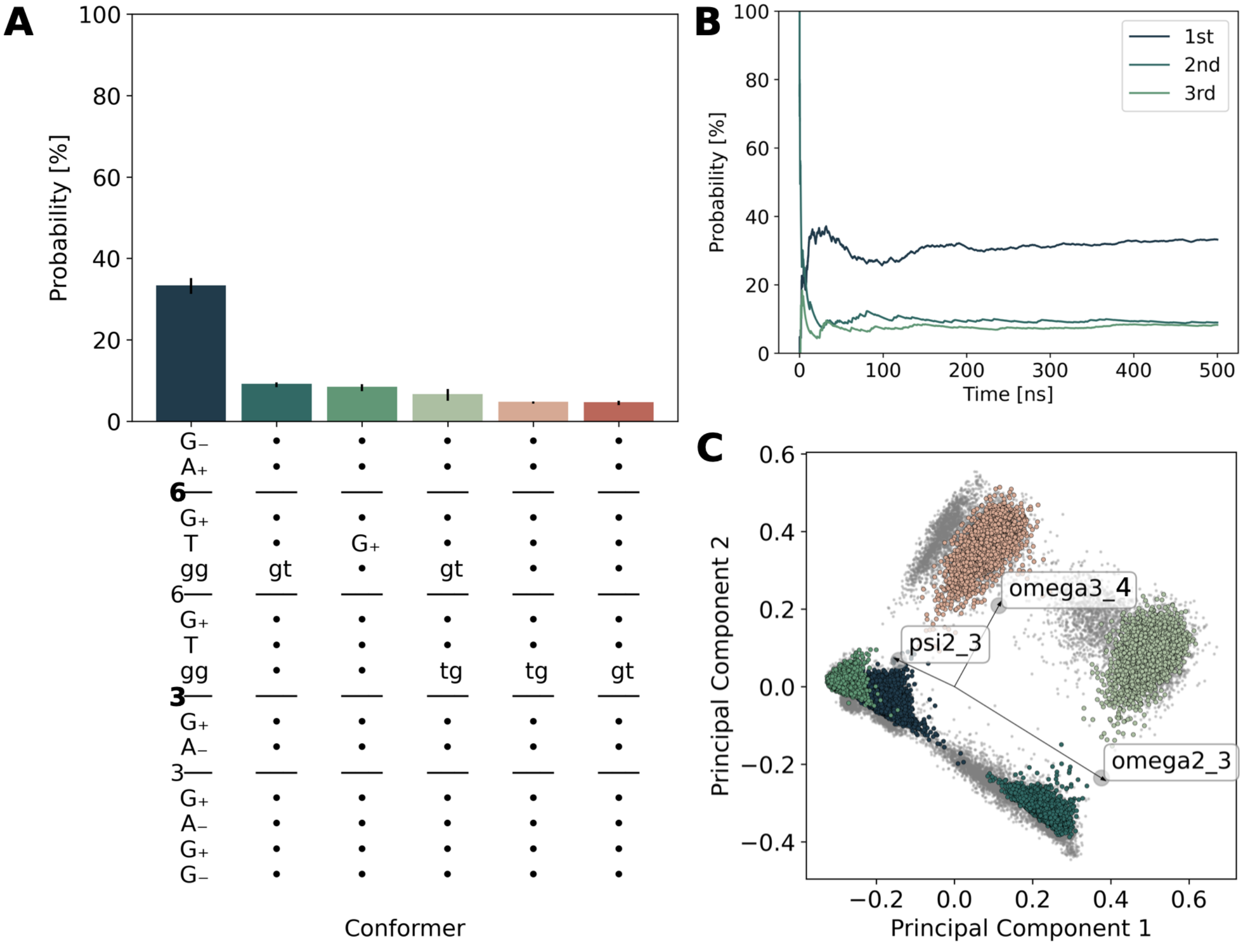


Figure S1: GlyCONFORMER[S1] analysis of M5G0 in solution displaying the conformer distribution as a histogram with each conformer representing one bin **A**, the cumulative average of the three most dominant conformers **B** and the conformational phase space of **A** represented in two dimensions via PCA, using all dihedral angles as input features. Vectors indicate the original feature axes with highest variance, where they point in the direction with highest squared multiple correlation with the principle components.

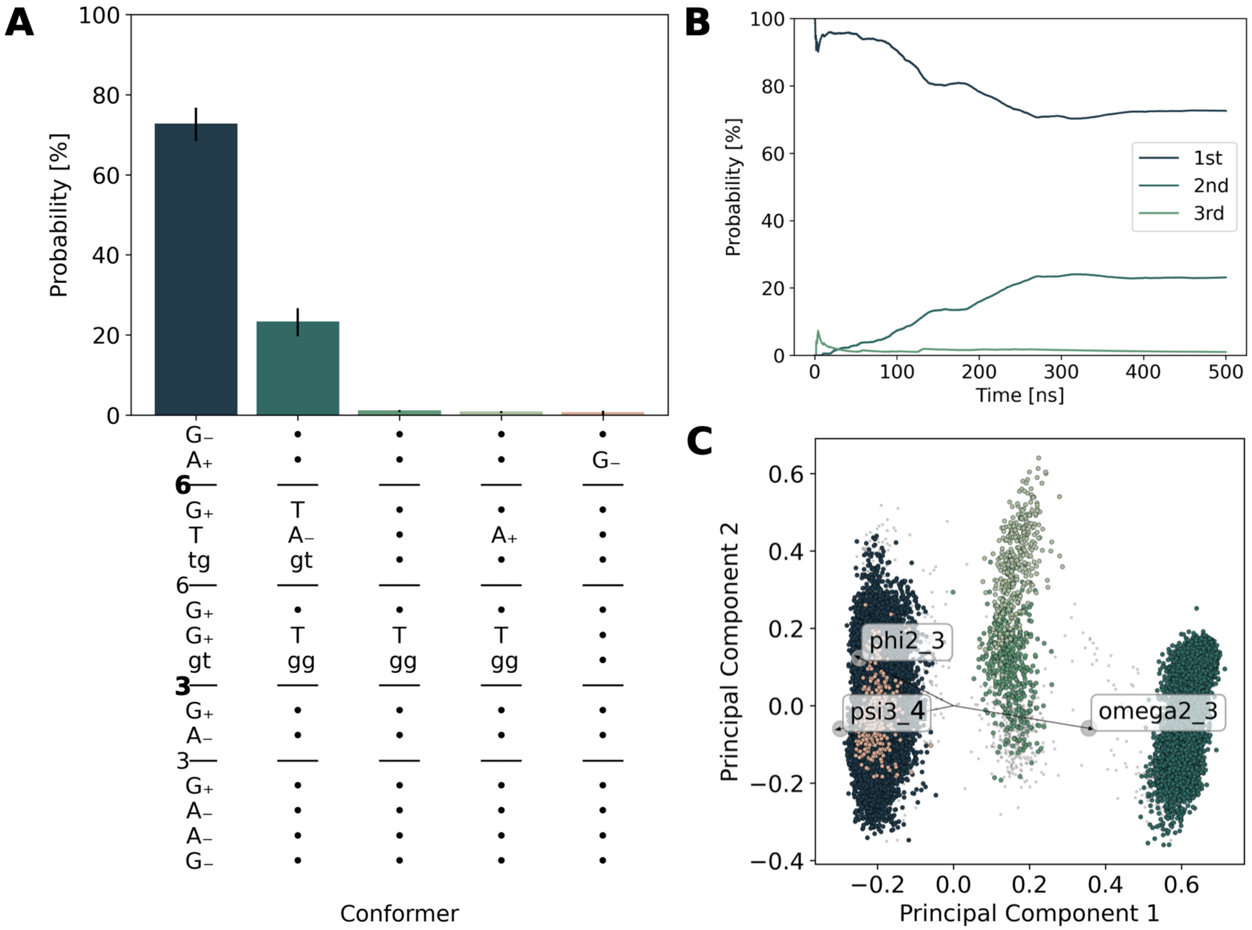


Figure S2: GlyCONFORMER analysis of M5G0 bound to MII with same panels as in Figure S1.

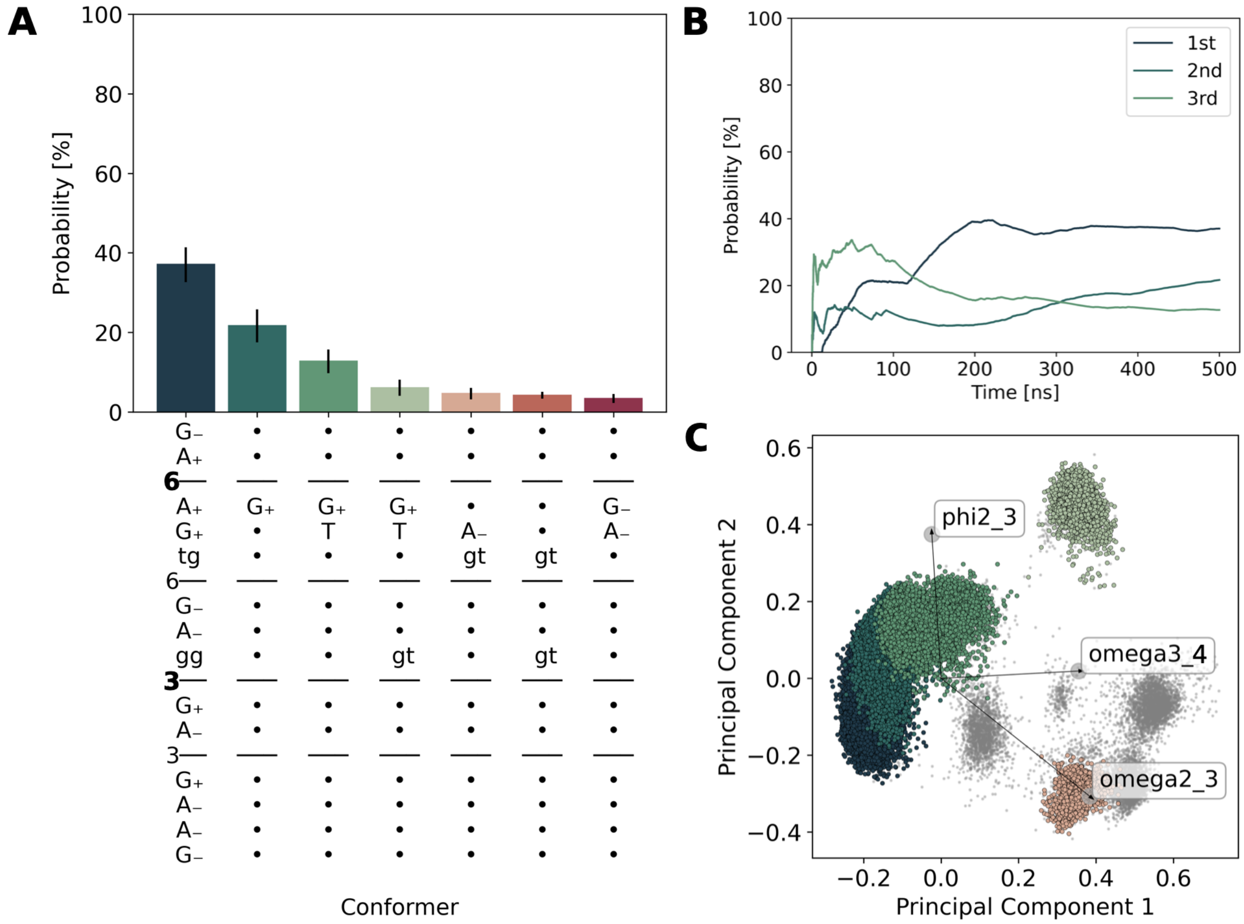


Figure S3: GlyCONFORMER analysis of M5G0 bound to MII with a restrain on the $H_{D341} - O6$ distance between amino acids D341 and the oxygen of the glycosidic linkage to be cleaved. Same panels as in Figure S1.

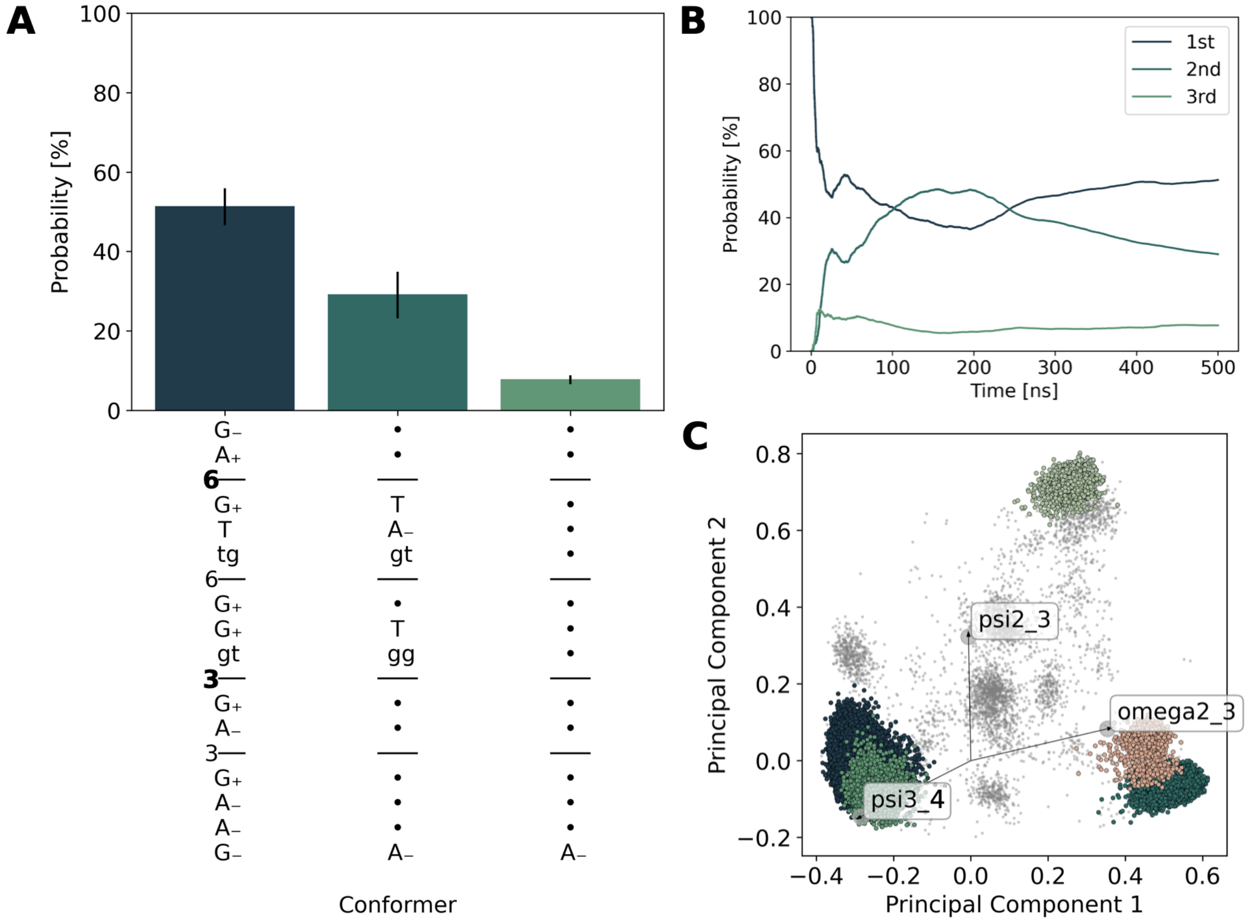


Figure S4: GlyCONFORMER analysis of M5G0 bound to mutant D204A MII with same panels as in Figure S1.

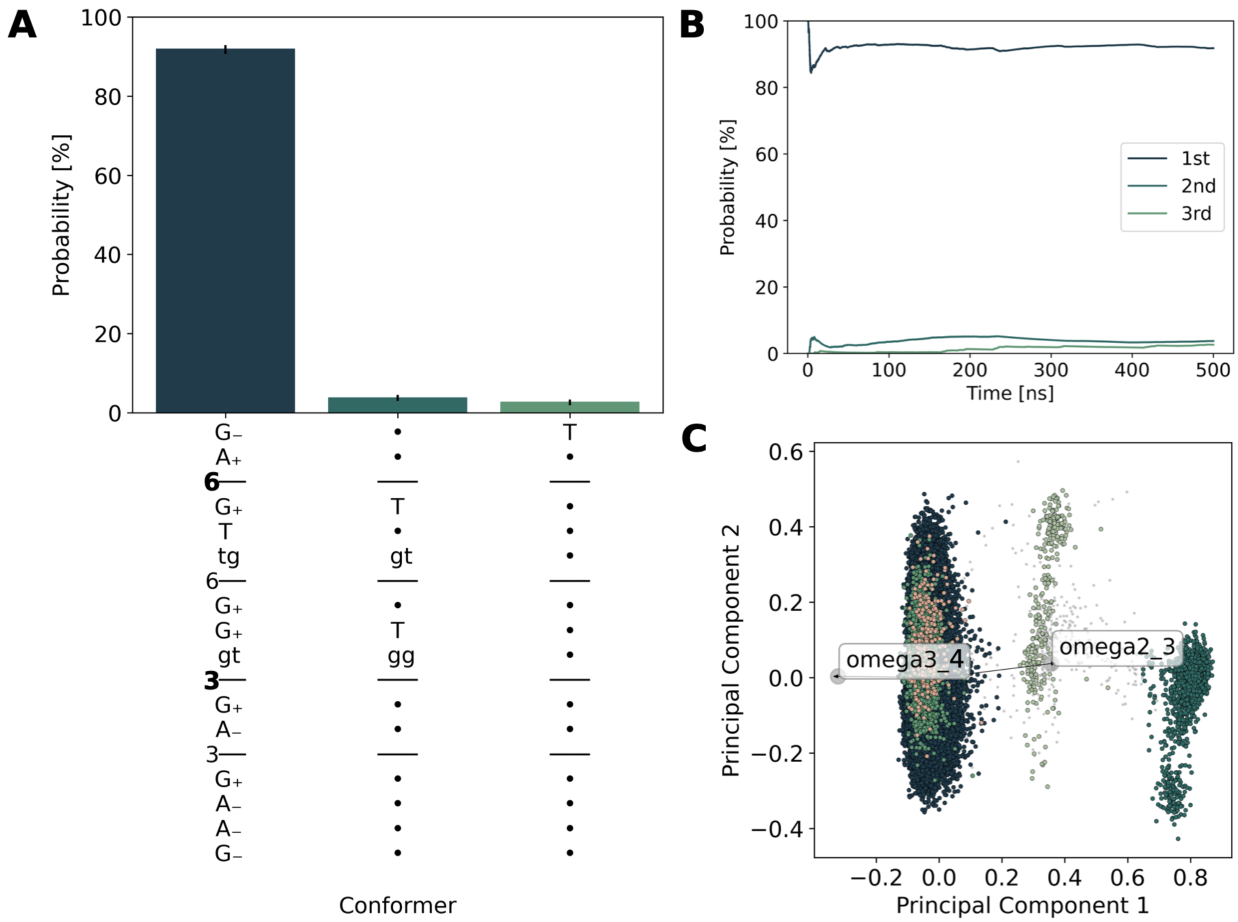


Figure S5: GlyCONFORMER analysis of M5G0 bound to mutant D341A MII with same panels as in Figure S1.

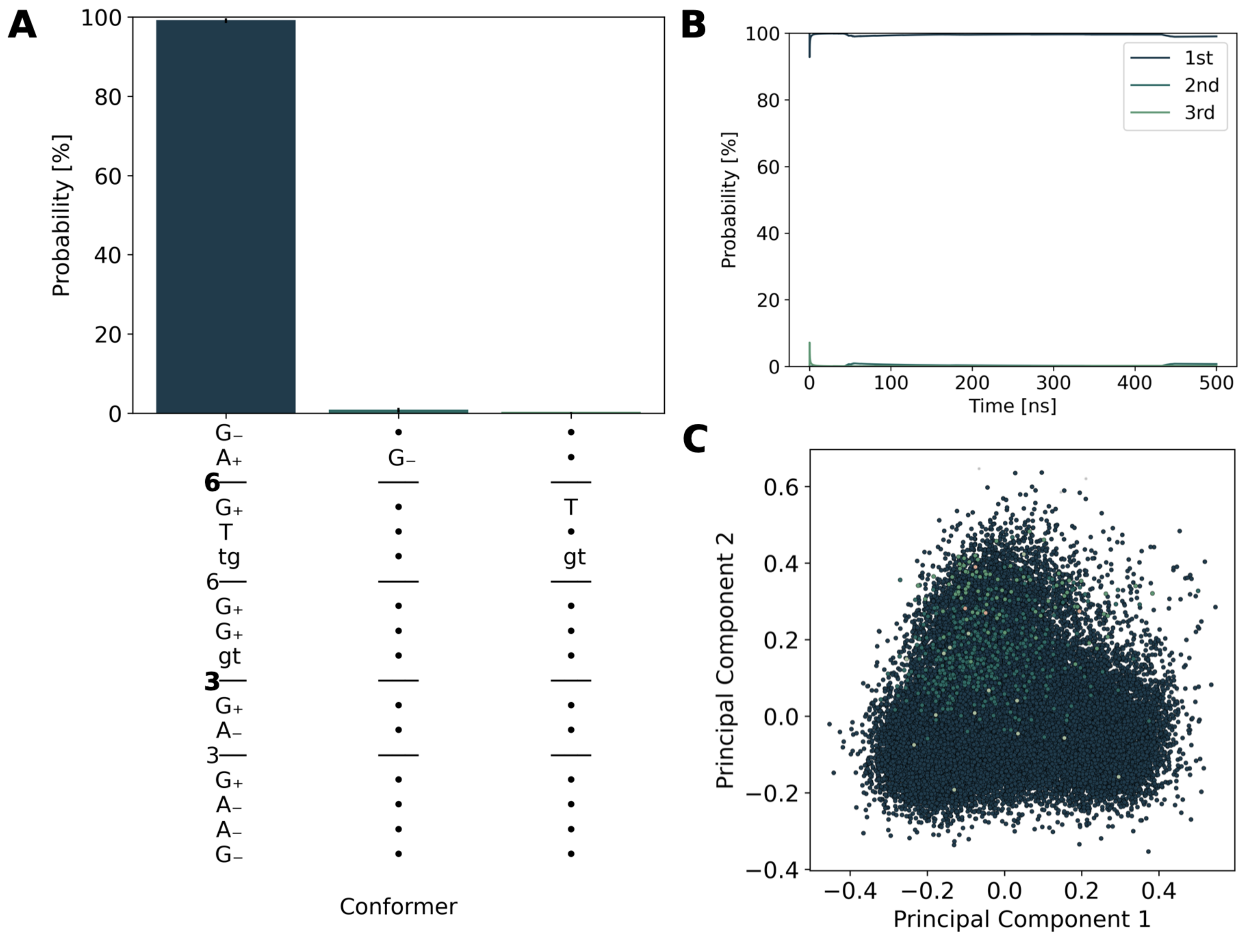


Figure S6: GlyCONFORMER analysis of M5G0 bound to mutant D92A MII with same panels as in Figure S1.

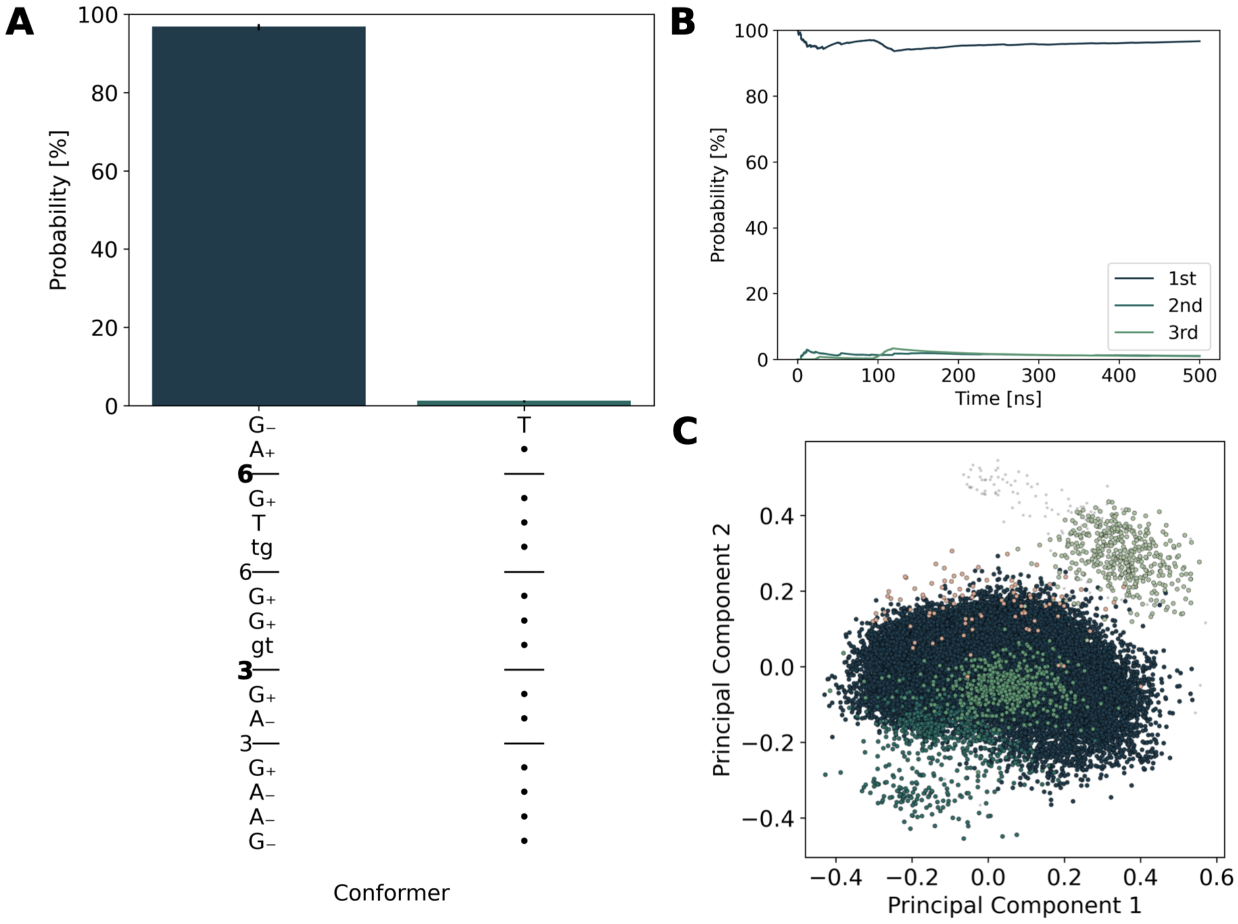


Figure S7: GlyCONFORMER analysis of M5G0 bound to mutant D472A MII with same panels as in Figure S1.

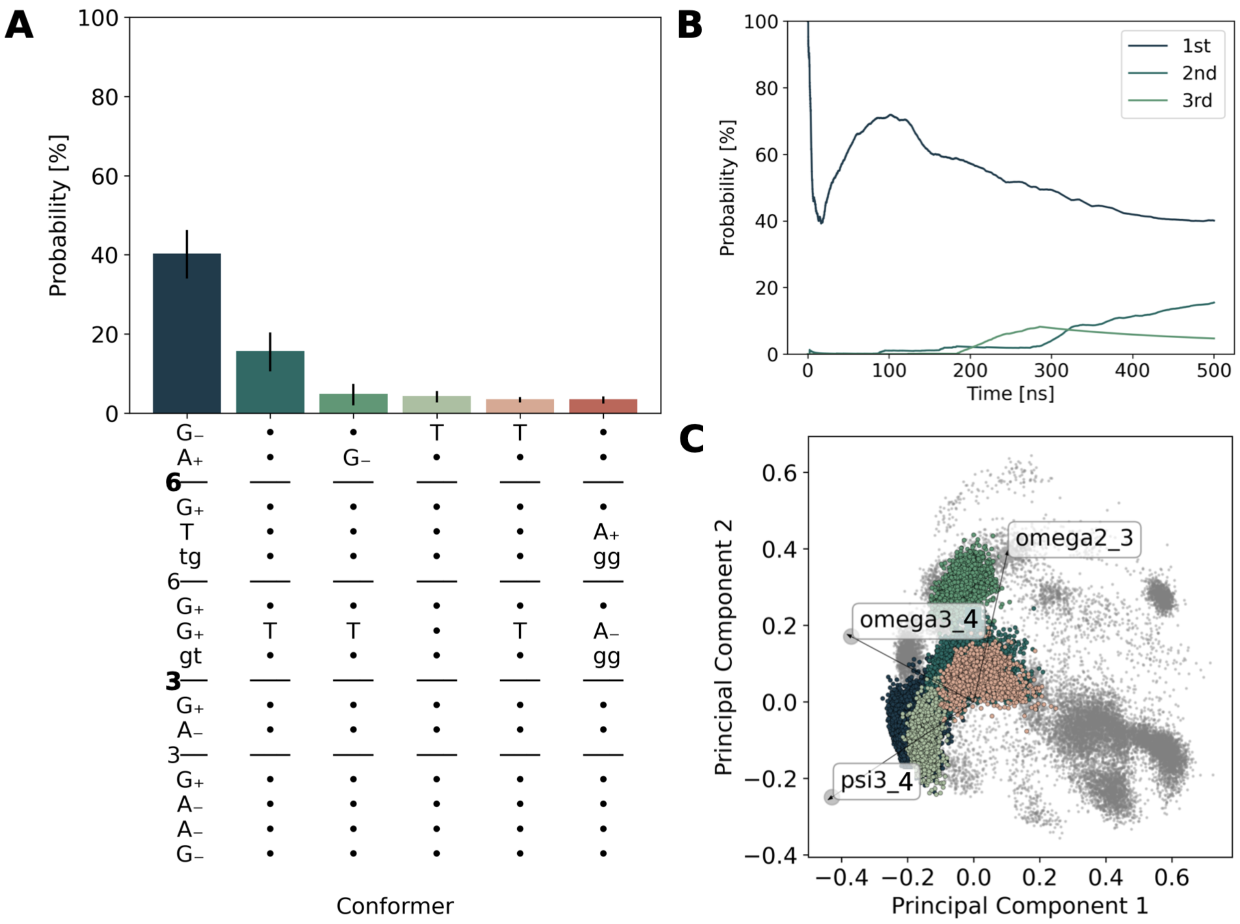


Figure S8: GlyCONFORMER analysis of M5G0 bound to MII lacking the Zn ion with same panels as in Figure S1.

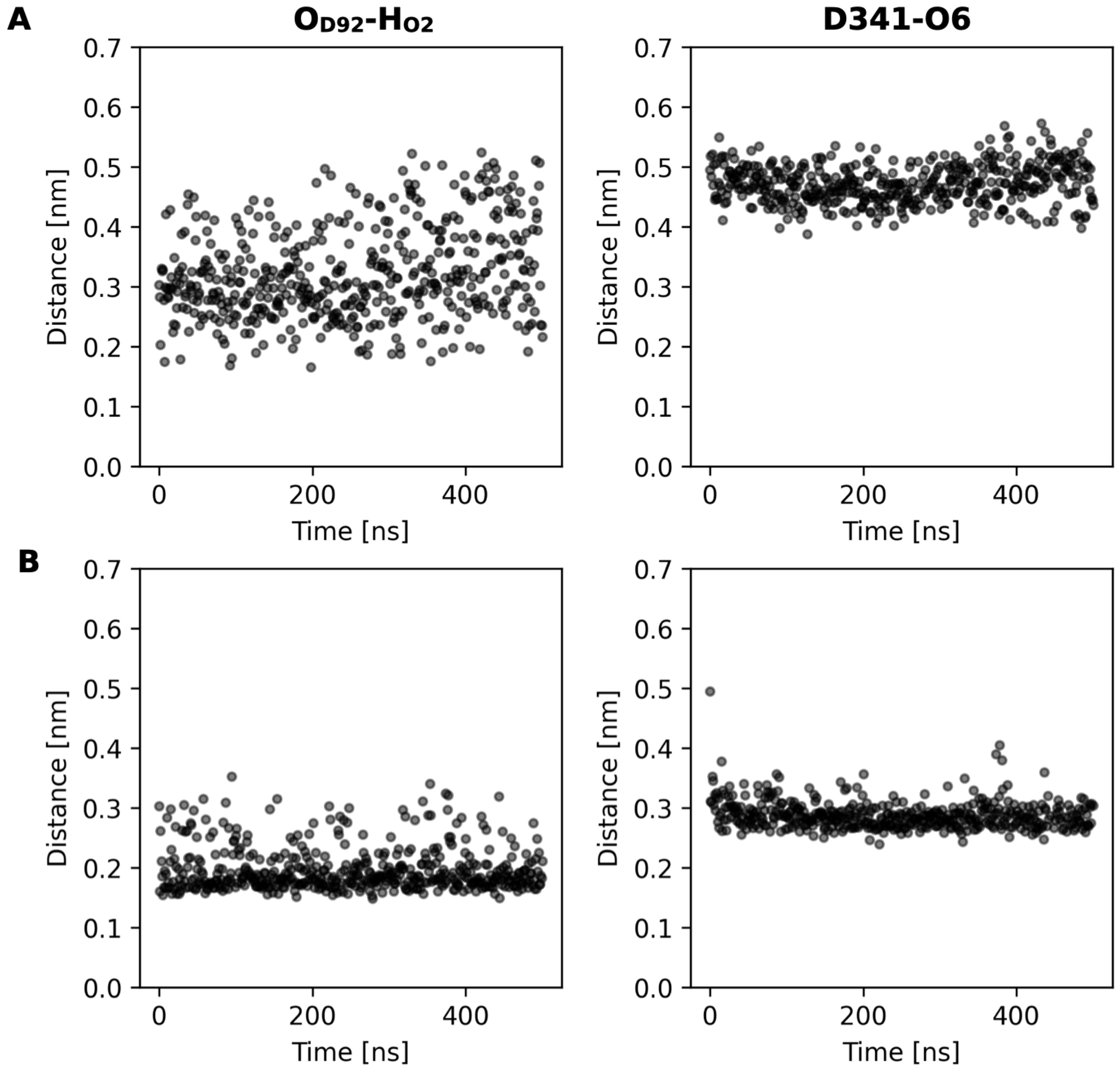


Figure S9: Distances between amino acid and glycan atoms that are critical for ring distortion captured for **A** M5G0+MII and **B** M5G0+MII with the $H_{D341}$-O6 restrain over time, plotting the behavior in the ground replica.

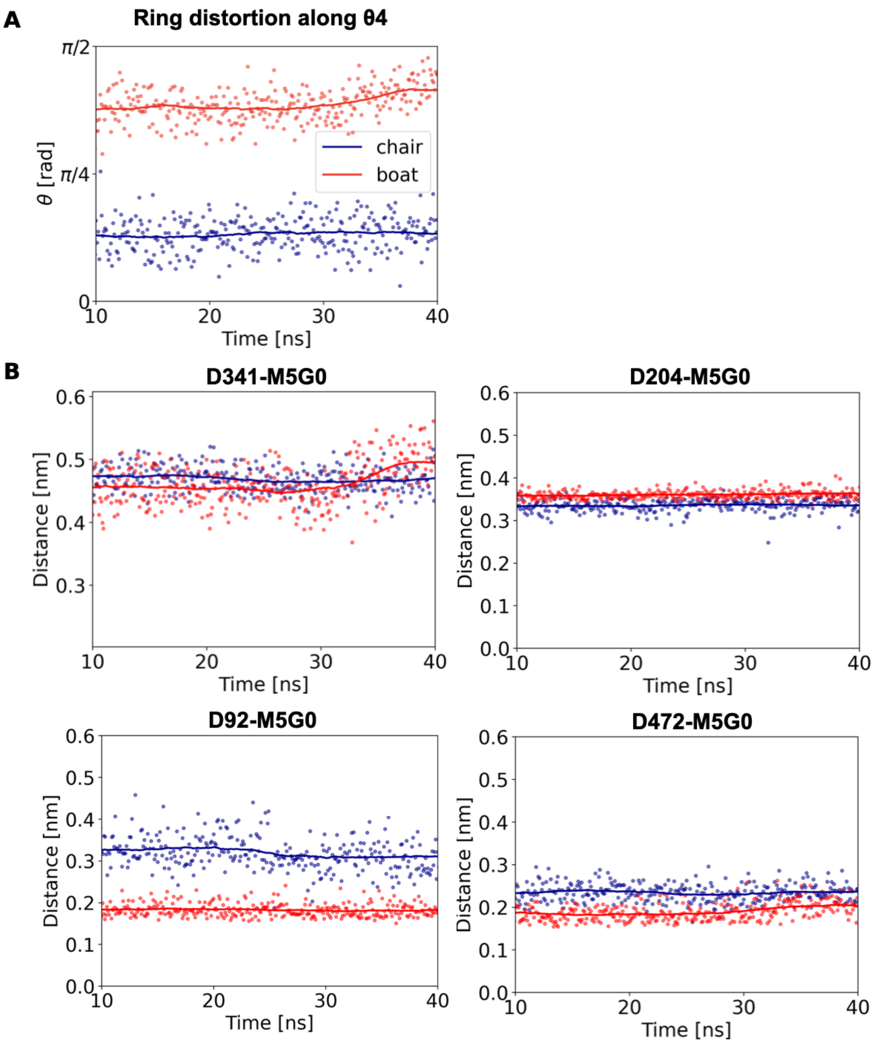


Figure S10: Distances between certain amino acids and the bound glycan M5G0 recorded from **A** two classical MD simulations where Man4 is positioned in the $^4C_1$ chair (blue) or $^0H_5$ half-chair (red). The chair was the equilibrium structure derived from the crystal structure. The half-chair was induced by restraining the distances $O_{D92} - H_{O2}$ and $H_{D341} - O6$ to 0.15 nm for 50 ns prior to sampling under unrestraint conditions. **B** Specific distances are $H_{D341} - O6$ (D341-M5G0), $O1_{D204} - H_{O2}$ (D204-M5G0), $O2_{D92} - H_{O2}$ (D92-M5G0) and $O1_{D472} - H_{O4}$ (D472-M5G0). Dots represent individual data points and lines the moving average with a window size of 100 data points.

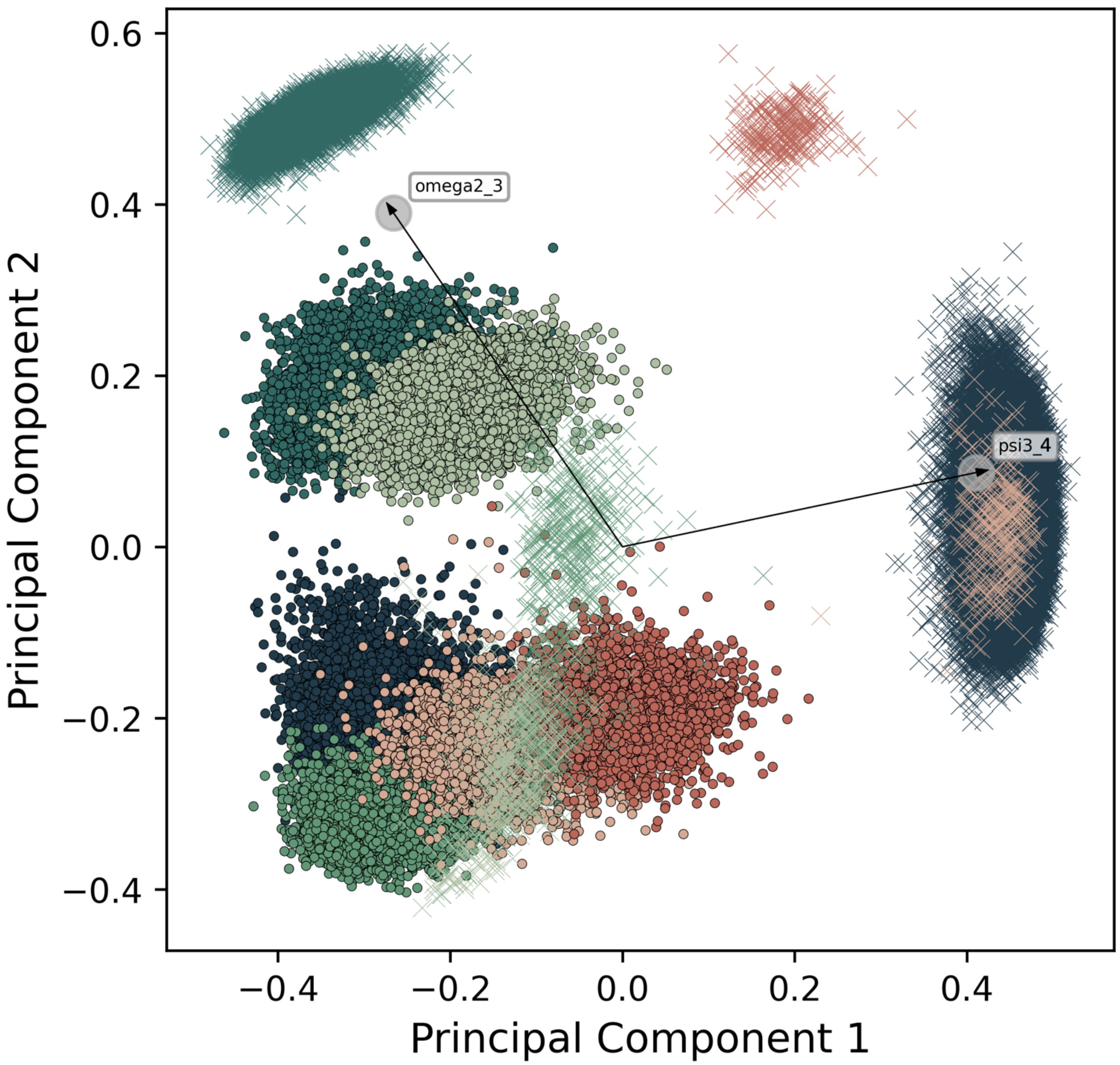


Figure S11: Comparative conformational phase space for M5G0 in solution (dots) and bound to MII (crosses) projected along PC1 and PC2. Only data points corresponding to the 6 most prominent conformers are plotted. Vectors indicate the original feature axes with highest variance, where they point in the direction with highest squared multiple correlation with the principle components. Colors correspond to the conformers and match the once in Figure S1 and S2.

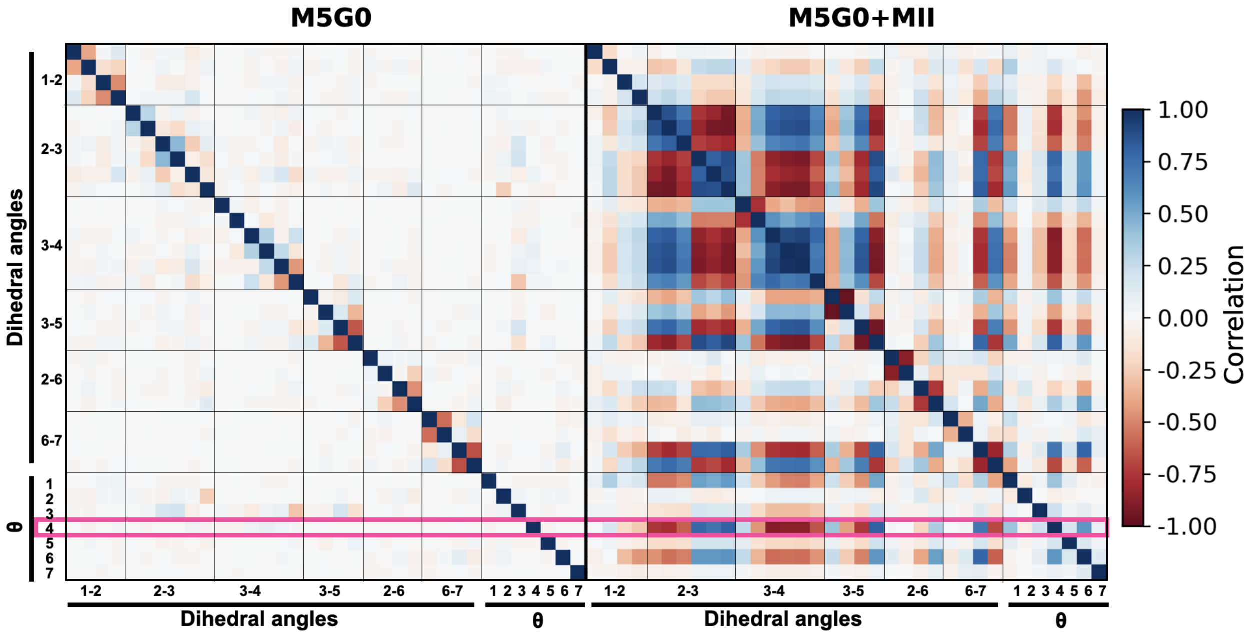


Figure S12: Correlation matrices for M5G0 in solution (left) and bound to MII (right), displaying the Pearson correlation coefficient for all dihedral angles (sin and cos) and $\theta$ of all monosaccharides. The pink square highlights the ring distortion of Man4.

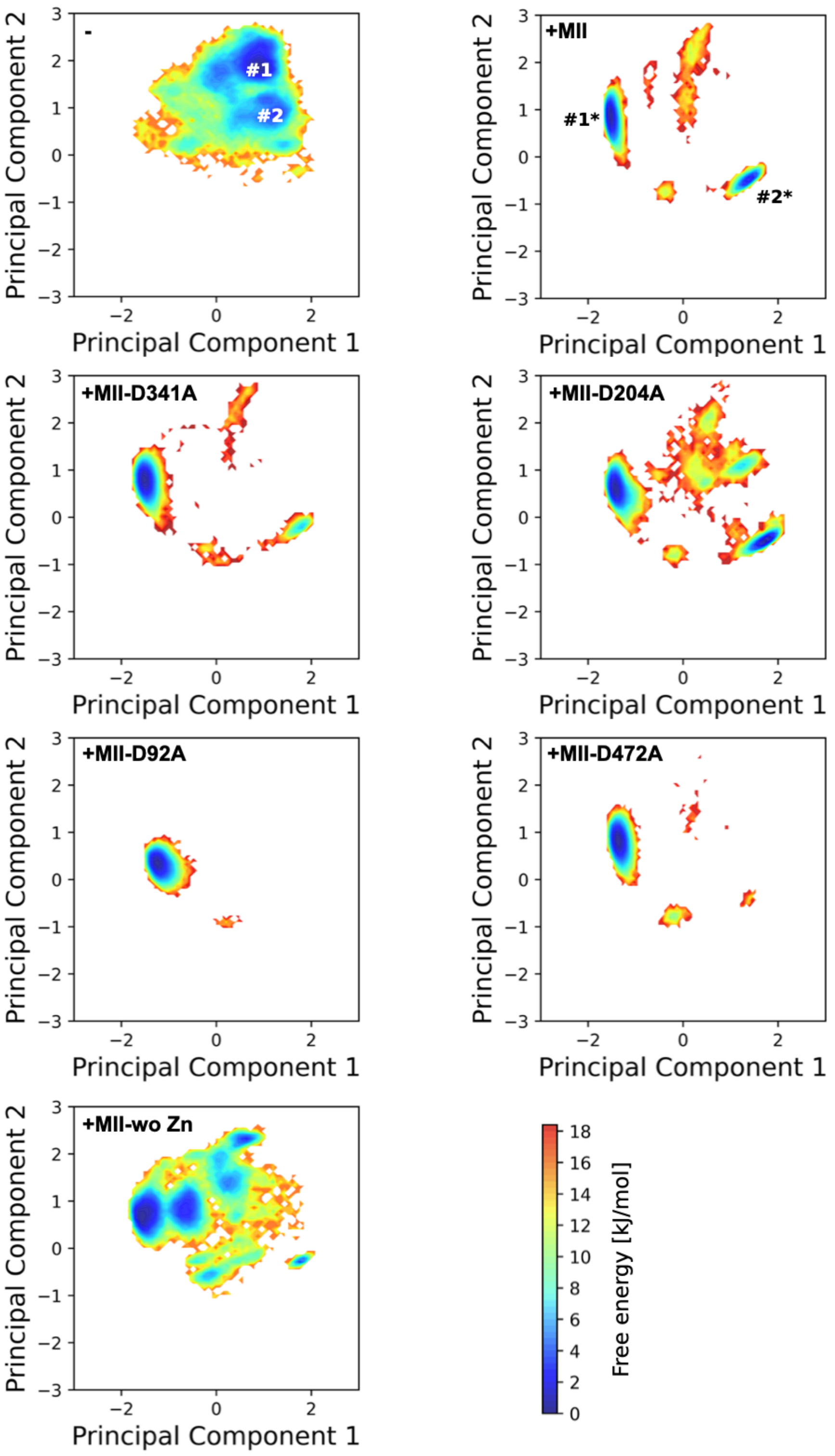


Figure S13: PCA of the comparative conformational phase space for M5G0, M5G0+MII or a mutated variant. First two glycan conformers for M5G0 and M5G0+MII are indicated with labels.

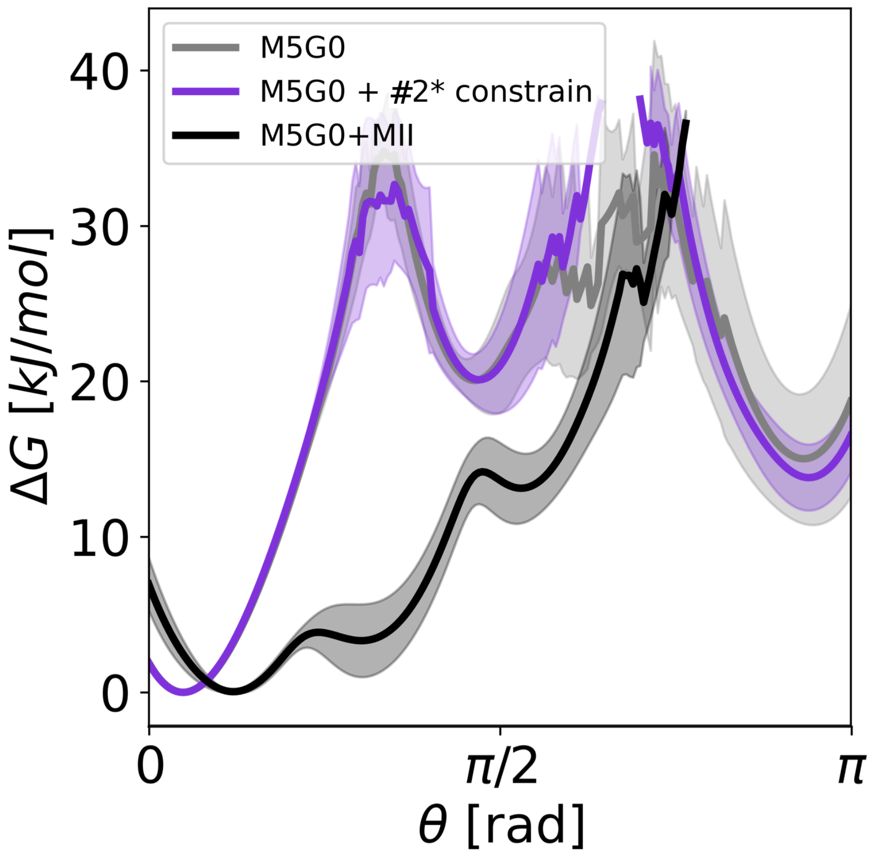


Figure S14: Ring distortion of the terminal mannose residue in glycan M5G0 monitored by the 1D Cremer-Pople parameter $\theta$ for M5G0 in aqueous solution (- enzyme), MII's catalytic site at subsite *-1* (+ enzyme) and the conformation M5G0 restricted to the #2* conformer, corresponding structurally to the enzyme bound state that has a distorted ring.

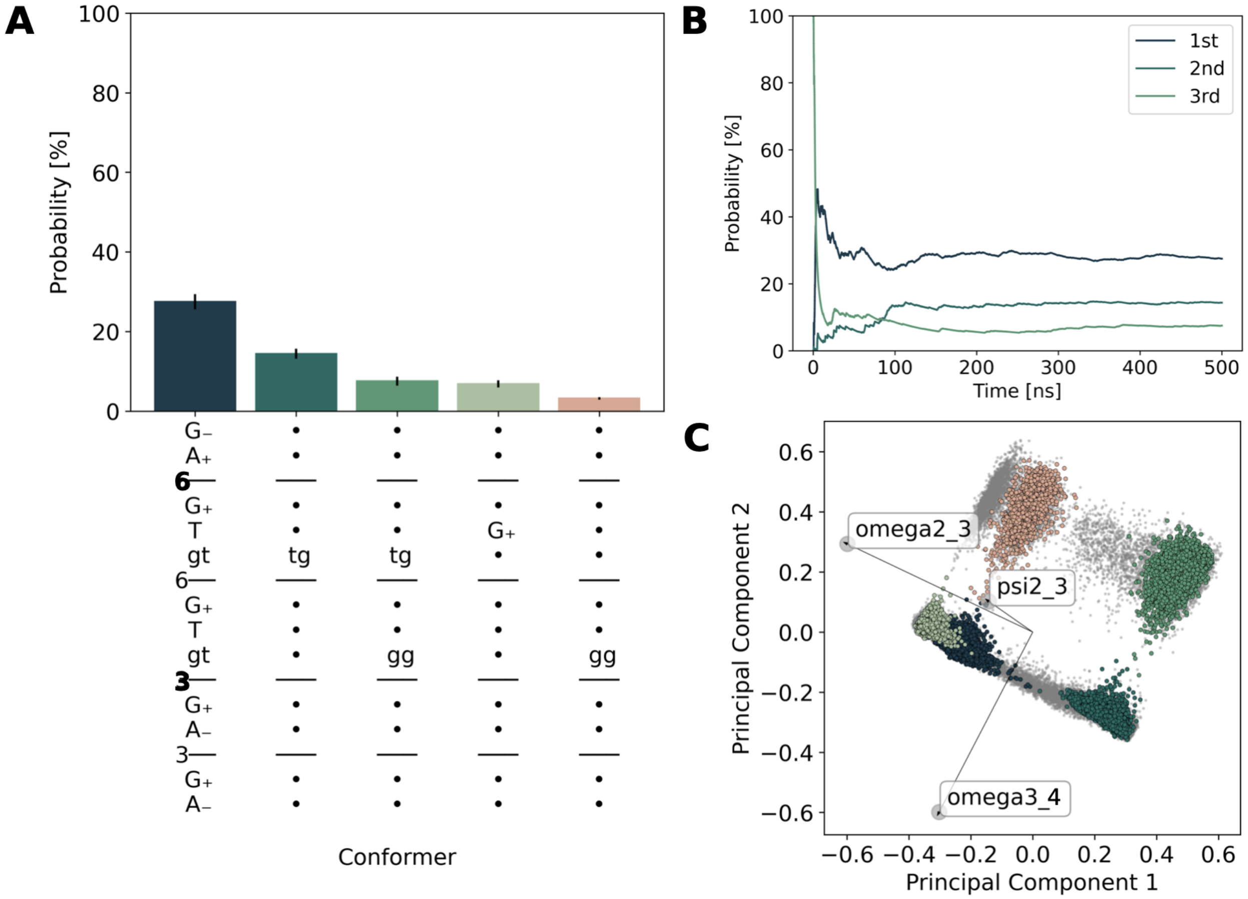


Figure S15: GlyCONFORMER analysis of M5 in solution with same panels as in Figure S1.

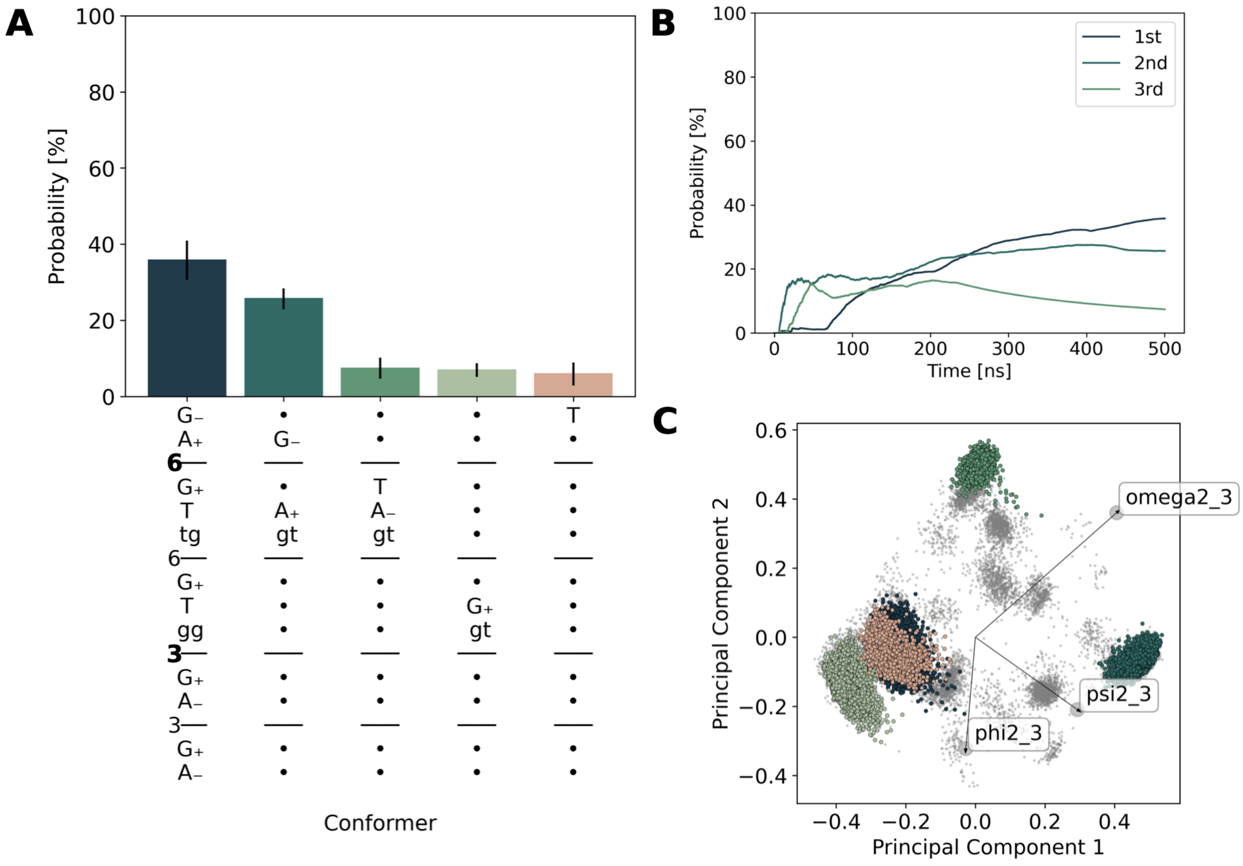


Figure S16: GlyCONFORMER analysis of M5 bound to MII with same panels as in Figure S1.

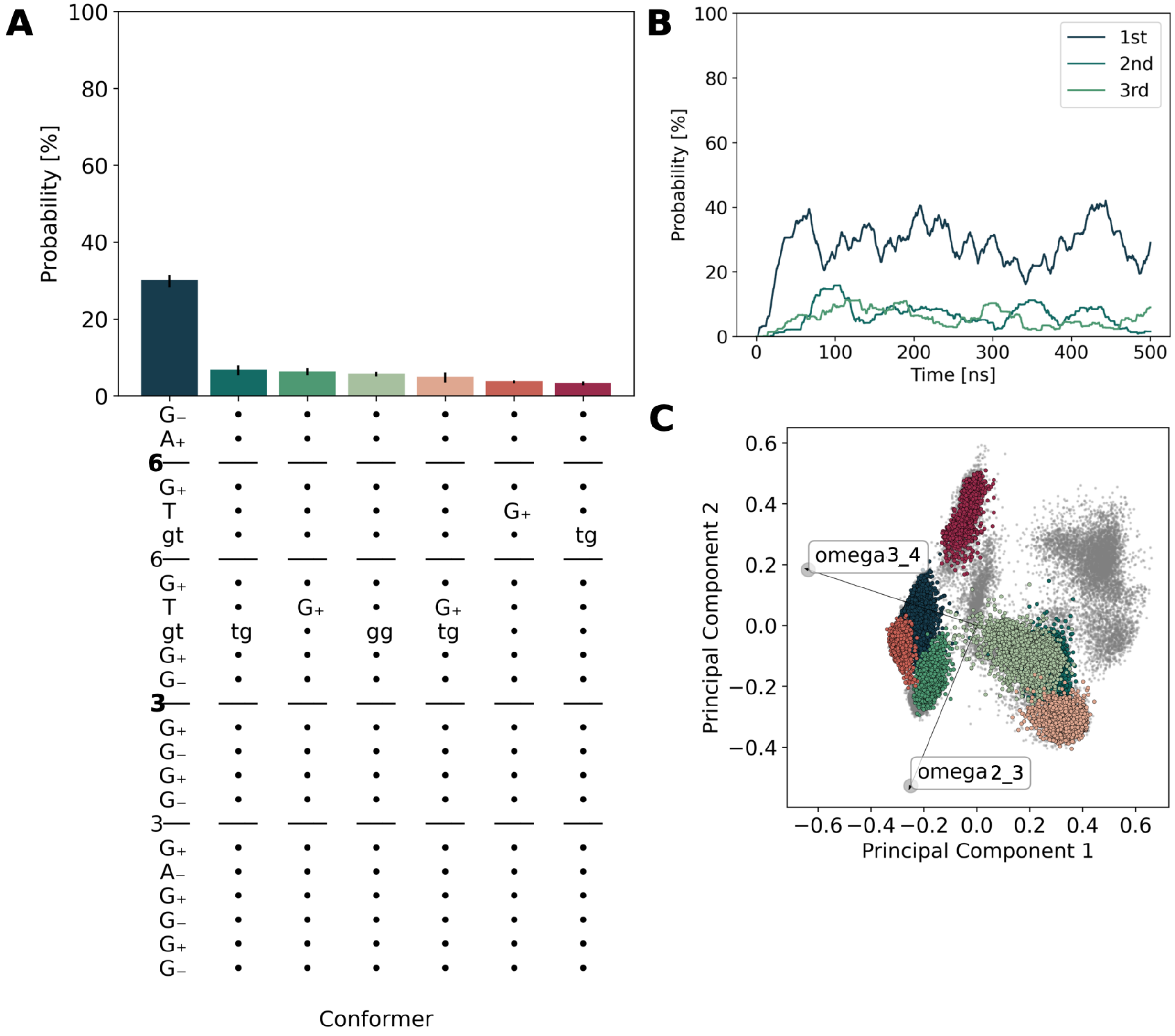


Figure S17: GlyCONFORMER analysis of M9 in solution with same panels as in Figure S1.

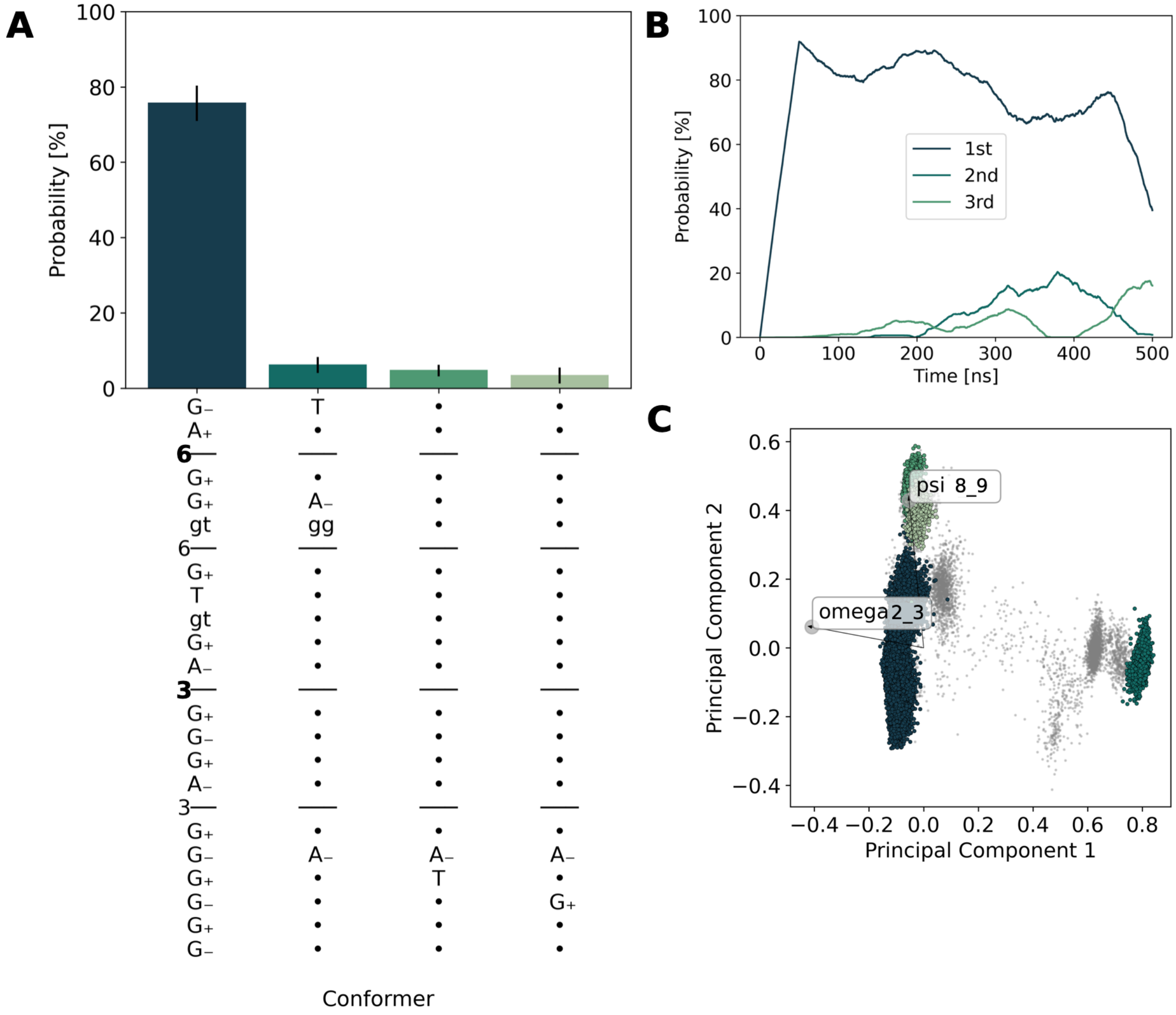


Figure S18: GlyCONFORMER analysis of M9 bound to MI (M9+MI) with same panels as in Figure S1.

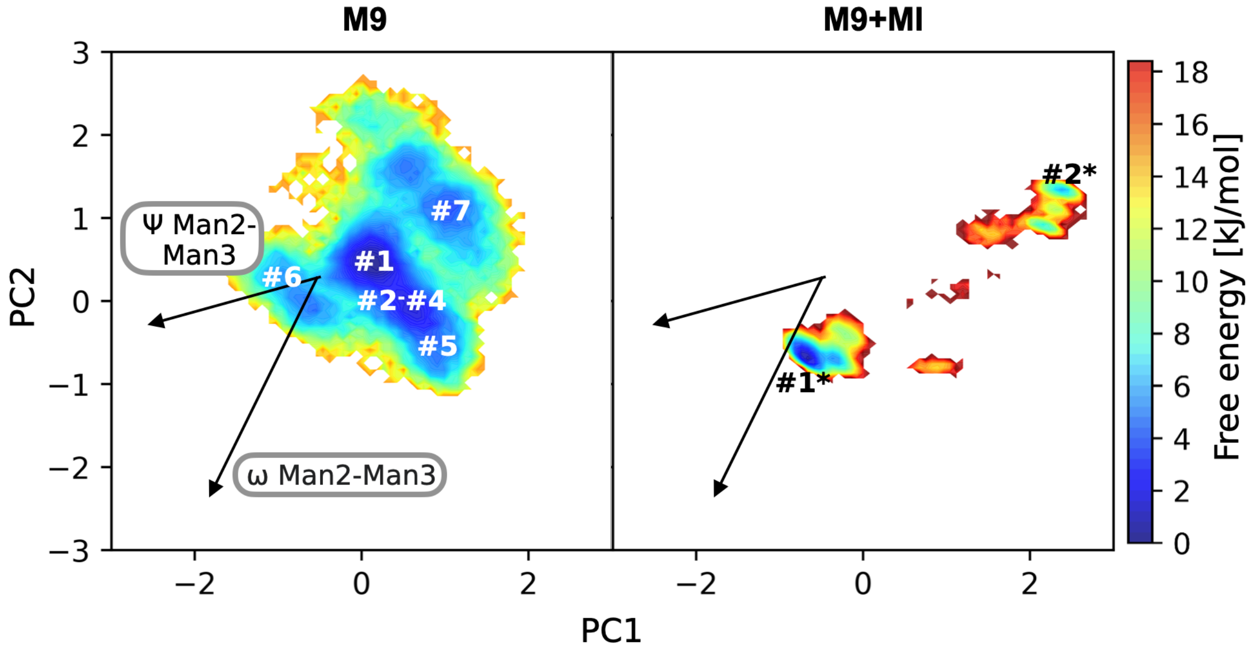


Figure S19: Comparative free energy surfaces of the conformational phase space for M9 and M9+MI projected along PC1 and PC2, with labeled conformers and vectors indicate the original feature axes with highest variance.
